# Modulation of motor cortical theta and gamma oscillations using phase-targeted, closed-loop optogenetic stimulation of local excitatory and inhibitory neurons

**DOI:** 10.1101/2025.08.21.670039

**Authors:** Jessica Myatt, Robert Toth, Timothy Denison, Isaac Grennan, Colin G McNamara, Charlotte J Stagg, Andrew Sharott

## Abstract

Theta and gamma oscillations are prominent features of cortical local field potentials (LFPs) and stimulation of the motor cortex at these frequencies can enhance motor learning. Phase-targeted closed-loop stimulation could provide a more precise and effective method to modulate these oscillations, particularly if stimulation parameters could harness the dynamics of the specific circuit mechanisms underpinning the generation of these activities.

To address this question, we defined the response of theta-and gamma-frequency oscillations in the motor cortex to closed-loop optogenetic stimulation of excitatory pyramidal neurons and inhibitory interneurons transfected with Channelrhodopsin-2 in awake, head-fixed RBP4-Cre (retinol-binding-protein-4) and PV-Cre (parvalbumin) mice, respectively. Phase-targeted blue-light pulses were delivered using the OscillTrack algorithm to track theta phase in the cortical LFP in real time and trigger stimulation at one of four target theta phases. Stimulation was delivered over a quarter of the target theta cycle, either as a single continuous pulse (“continuous” protocol) or three short pulses at gamma (75Hz) frequency (“gamma” protocol).

Stimulation of both neuron types, using either stimulation protocol, modulated theta power in a phase-dependent manner, with continuous stimulation of excitatory cells leading to stronger modulation. Phase-dependent amplification during stimulation of excitatory vs inhibitory neurons was offset by 90°, in line with predictions from computational models. Open-loop replay of previously recorded closed-loop stimulation patterns did not elicit the same phase-specific effects, demonstrating the necessity of the closed-loop interaction to produce these effects. Stimulation of pyramidal neurons using the gamma protocol amplified gamma power, independently of target theta phase.

These findings reveal phase-dependent amplification of cortical theta power can be induced by stimulation of local excitatory or inhibitory neurons, with a phase-offset likely resulting from circuit interactions. This approach can be used to inform the development of brain stimulation methods to modulate these activities more effectively in humans.

## Introduction

Oscillatory activity in the cortex emerges from the dynamic interplay between excitatory pyramidal neurons and inhibitory interneurons, which produce a fluctuating excitatory-inhibitory balance and tightly regulates the spiking activity within cortical circuits [1–4]. These processes play key roles in orchestrating information processing, communication and coordination across large neural networks, with roles in cognition, perception, memory and motor control. Moreover, abnormalities in oscillatory activities have been identified in several neurological and neuropsychiatric disorders [5]. Interventions that can correct these changes could provide a powerful approach to treatment.

In the motor cortex, theta and gamma frequency oscillations may have specific roles in motor adaptation and learning [6]. In line with this, theta-gamma coupled transcranial alternating current stimulation (tACS) of motor cortex can improve motor learning [7], suggesting that manipulating these activities could provide a powerful therapeutic tool. Such stimulation, however, does not take the intrinsic system dynamics into account. A more effective approach could be to couple the timing of stimulation to ongoing theta and/or gamma oscillations. Phase-targeted stimulation, where the phase of an ongoing oscillation is tracked in real time and the detection of a “target” phase is used to trigger stimulation, provides a precise and effective method to achieve this. Our previous studies, where electrical stimulation was delivered to specific phases of Parkinsonian beta [8] and hippocampal theta [9] oscillations, demonstrate the capability of this approach to both amplify and suppress the target activity. Such phase-targeted stimulation could be further improved by understanding and utilising local circuit dynamics. This is particularly relevant for cortical circuits, where the mechanisms underlying oscillations are often a function of interactions between diverse neuron types in the local microcircuit [10]. To this end, several studies have utilised phase-targeted optogenetic stimulation of specific cell types [11–16], but have not compared the effects of the same closed-loop protocol to stimulate different neuron types in the target circuit. Applying such an approach to motor cortex could provide insights into the general mechanisms underlying phase-targeted neuromodulation of motor cortical circuits and motor control, including which circuit components are the key drivers. Given the ubiquitous presence of theta and gamma oscillations across cortical circuits [6,17,18], developing closed-loop stimulation protocols targeting these oscillations may provide powerful tools for modulating a wide range of cortical functions.

In this study, we applied the same phase-targeted closed-loop stimulation protocols to two neuron types in the motor cortex of awake, head-fixed mice. Optogenetic excitation using Channelrhodopsin-2, was applied to excitatory pyramidal neurons in Layer V and parvalbumin-expressing (PV) inhibitory interneurons, using transfection in RBP4-Cre (retinol-binding-protein-4) and PV-Cre mice respectively. Stimulation was delivered at one of four target phases of the ongoing theta oscillation using two protocols; either a continuous pulse or a burst of gamma frequency stimulation. Our results reveal that stimulation of both cell types using either protocol results in phase-dependent modulation of the amplitude of ongoing motor cortical theta oscillations, but that the amplifying phase differs across the neuron types. We further demonstrate that stimulation of excitatory neurons may more effectively evoke changes in gamma power. These findings provide new insights into circuit-level mechanisms shaping motor cortical oscillations, thereby informing strategies for more effective targeted therapeutic neuromodulation in using non-invasive or deep brain stimulation approaches.

## Materials & Methods

### Animals

All studies were performed in accordance with institutional guidelines and the Animal (Scientific Procedures) Act, 1986 (United Kingdom). Experimental procedures were conducted in adult mice (male) either B6;129P2-Pvalb^tm1(cre)Arbr^ (PV-Cre, heterozygous on a C57BL/BJ background (n=11), Jackson Laboratories) or Tg(RBP4-Cre)KL100Gsat (RBP4-Cre, heterozygous with a C57BL/BJ background, Jackson Laboratories) (n=14), weighing 23-43g. Additional mice (n=2 PV-Cre; n=1 RBP4-Cre) were used for verification of the immunohistochemistry protocol. Mice were kept in a 12-hour light/dark cycle, with food and water provided *ad libitum*.

### Surgical procedures

All surgical procedures were performed under gaseous isoflurane in oxygen anaesthesia (4% for induction, maintained at 1-2%). Anaesthetised mice were secured in stereotaxic apparatus (David Kopf Instruments), and body temperature was maintained at 37±0.5°C. Analgesia (Buprenorphine; 0.08mg/kg in sterile saline) was administered subcutaneously after induction of anaesthesia. Prior to viral injection and headplate surgeries, a local anaesthetic (Marcain; 2mg/kg) was injected into the scalp subcutaneously. Following surgical procedures, a subcutaneous injection of 0.4ml pre-warmed isotonic glucose solution was given.

### Viral injection

Viral expression in neurons was achieved by unilaterally injecting 100nl of AAV5-Ef1a-DIO-ChR2(E123T/T259C)-eYFP (∼6.2×10^12^ vg/ml; Deisseroth Vector Core, University of North Carolina) (ChR2) or control AAV5-Ef1a-DIO-eYFP (∼4×10^12^ vg/ml; Deisseroth Vector Core) vector (which will not express ChR2) into the right motor cortex (AP +1.5mm, ML +1.3mm, DV +0.8mm; from Bregma).

### Headplate implantation

Mice were implanted with a titanium custom-made headplate (Get It Made Ltd, London) (0.7g, internal diameter 9mm) to enable head-fixation. Skin overlying the skull was removed, the skull was washed (3%, H_2_0_2_ in dH_2_0 followed by sterile saline), then etched for improved adhesion. Edges of skin were glued (Vetbond) to the skull, before the headplate was aligned to bregma and lambda, then bonded to the skull using clear dental cement (Super Bond C&B, Parkell), a thin layer of which was also used over the entire exposed skull surface. A craniotomy was made to insert a reference screw (Precision technology, East Grinstead) wrapped in wire over the ipsilateral cerebellum to the injection site, this was fixed with dental cement (Jet Denture Repair Power: Lang Dental, Illinois, USA; Meadway Repair Liquid: MR. Dental, Surrey).

### Craniotomy for electrophysiological recordings

The day prior to recording, a small craniotomy (∼1.5mm^2^) was made above the recording site. To provide a protective barrier over exposed brain tissue, dura gel (Cambridge NeuroTech) and silicone (Body Double, Smooth-On) were used following the surgery and between electrophysiological recordings.

### Electrophysiological recordings

Electrophysiological recordings (maximum 5 per mouse) were undertaken in awake head-fixed mice previously habituated to the rig, in which the mouse sat head-fixed in a cardboard tube, with a cork lined 3D printed insert, supported by a caddy. A faraday cage was used to shield the set-up from electromagnetic signals in the external environment.

To record local field potentials in the motor cortex (1000-1300mm in depth), linear four shank 32 channel probes (100mm between contacts, 200mm between shanks; A4×8-100-200-177-OA32; NeuroNexus, Michigan, USA) were used for acute recordings. A cortical headstage (64-channel; Intan Technologies, California, USA) was attached to the probe using an adaptor (Adpt-A32-OM32, Omnetics, Minnesota, USA) and connector (OM32, NeuroNexus), as well as to both a grounding wire and cerebellar reference screw. All electrophysiological recordings were collected using the Intan RHD2000 system at a sampling rate of 20kHz. The probe was positioned above the desired coordinates using a stereotaxic frame and arm (Kopf Instruments, California, USA). In later recordings, the probe had an additional ‘Z-Coat’ (NeuroNexus, Michigan, USA). On the second shank, 200mm above the top contact site was an integrated optic fiber, used to deliver optogenetic stimulation (473nm; Cobolt Laser, Sweden) via a connected optic fiber (105mm core, 200mm, 0.22NA, Thorlabs, New jersey, USA). Laser power was verified and adjusted for every session using a power meter interface (PM100USB, Thorlabs) and sphere photodiode power sensor (S140C, Thorlabs).

### Theta phase locked optogenetic stimulation

Online phase tracking of 6Hz motor cortical theta was performed using the OscillTrack algorithm [19] and used to drive optogenetic stimulation at specific phases of ongoing theta activity. The real-time phase estimate was calculated in the field-programmable gate array (FPGA) of the Intan RHD2000 USB interface board, allowing target phase and stimulation output to be selected on a GUI. In each recording session a phase reference channel was selected from visual observation of baseline spiking activity. This channel was high-pass filtered using a 1st order digital filter with a corner frequency of 4Hz to reduce the interference of intense 3Hz delta activity, then down-sampled 49-fold to a rate of 408Hz for processing. The algorithm was configured with a loop-gain of 0.0625 and a centre frequency of 6Hz in all sessions. A real-time trigger was only generated at the target phase if more than 80% of the period of the 6Hz theta cycle had passed (timeout: 133.333ms) to reduce erroneous triggering [8].

### Optogenetic stimulation protocols

Stimulation patterns (Table 1.1 & Fig. 1A-C) were delivered to the motor cortex for a period of ∼2.5-7 minutes over four different phases of the 6Hz theta cycle, centred to the target phase, in a randomised order at a combination of different laser powers (0mW (unstimulated control), 1mW, and 2mW). *Continuous* stimulation consisted of a single, uninterrupted pulse delivered for a duration of approximately 1/4 of the 6Hz theta cycle. *Gamma* stimulation consisted of 3 short pulses (each 6667µs) evenly spaced at a frequency equivalent to 75Hz (see Table 1.1). *Arrhythmic gamma* stimulation was used to interrogate whether gamma rhythmicity was necessary to produce effects seen in response to the gamma protocol. As such, the middle of the three pulses varied in position between the two outside pulses based on a set of 31 positions, leading to stimulation that delivered the same energy over the same overall time window, but without the periodic component of the main protocol.

**Figure 1:**
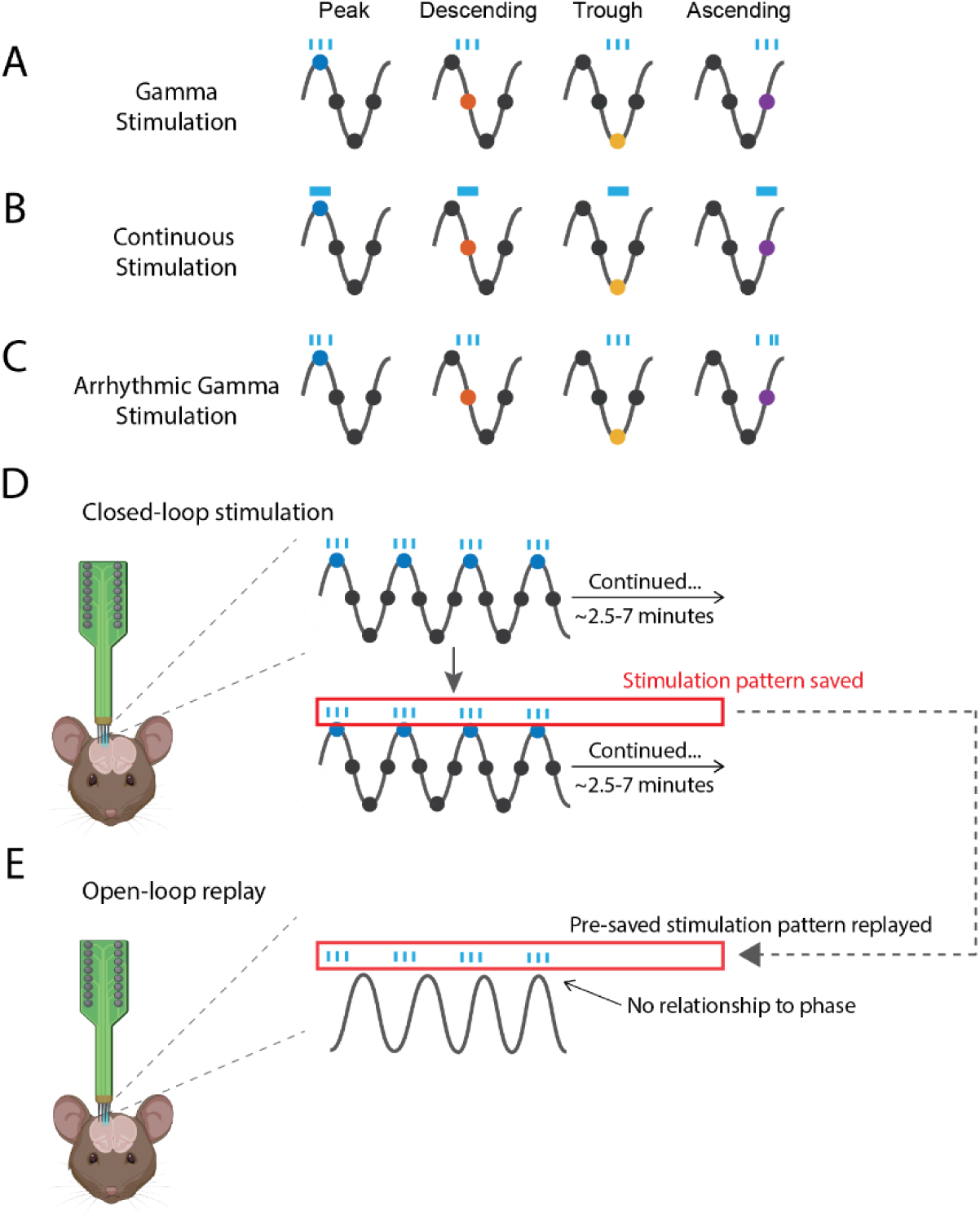
Stimulation patterns and targets for phase-targeted closed-loop and open-loop replay optogenetic stimulation. (**A** & **B**) Two optogenetic stimulation patterns were devised to interrogate whether there was differential modulation over 4 different phases of the theta cycle. The first was a burst of 3 pulses at gamma frequency (75Hz) (**A**), with the middle pulse centred to the phase, and the second a simple ON/OFF pattern over ¼ of the theta cycle (41.667ms) (**B**). In some experiments, an arrhythmic gamma protocol (**C**) was used to interrogate the importance of gamma rhythmicity in the circuit. In this protocol, the two outside pulses had the same interval as gamma stimulation, but the middle pulse was placed randomly between them resulting in 3 pulses that were spaced within the total duration of the main gamma protocol, but with random interstimulus intervals. For each recording within a session, a stimulation pattern, stimulation phase and stimulation on (1 or 2mW) or off (0mW) were chosen in a randomised order then stimulated and recorded between ∼2.5-7 minutes. (**D**) The 6Hz theta oscillation was tracked using the OscillTrack algorithm on a custom Intan RHD2000 USB interface, used for both stimulation and electrophysiological recordings. During each session all closed-loop stimulation patterns were recorded. In later sessions, these specific patterns were transferred to an SD card to replay them to the mouse later within the same session (‘open-loop replay’) (**E**). Care was taken to replay them at the correct stimulation amplitude as the original closed-loop recording (1 or 2mW). Thus, open-loop replay stimulation delivered the same number of pulses, over same total time period and used the same stimulation pattern at the same amplitude as closed-loop sessions, but with no relationship to the ongoing theta oscillation (Figure created with the help of BioRender.com).

**Table 1.1:** Stimulation patterns used for closed-loop optogenetic electrophysiology.

| Stim Pattern | Stim length (μs) | Pulse on (μs) | Pulse off (μs) | Timeout (μs) | Number of pulses |
| --- | --- | --- | --- | --- | --- |
| Gamma | 33335 | 6667 | 6667 | 133333 | 3 |
| Continuous pulse | 41667 | 41667 | 0 | 133333 | 1 |
| Arrhythmic gamma | 33335 | 6667 | Random interval | 133333 | 3 |

To differentiate the effect of theta phase locked stimulation from the temporal pattern of optogenetic stimulation trains generated through closed-loop recordings, an ‘open-loop replay’ paradigm was introduced to later recording sessions (Fig. 1D-E). These were pre-recorded stimulation patterns generated during closed-loop stimulation (details as noted above) replayed using the Intan digital output files during the same recording session. Thus, the pattern of stimulation was the same, but with no relationship to the phase of the ongoing theta oscillation. Files were replayed through an Arduino-compatible microcontroller (Teensy 3.6; PJRC Ltd, Sherwood, Oregon, USA) randomly, taking care to use the original laser amplitude, while triggering recording of electrophysiological data through Intan RHD2000.

## Data Analysis

Welch power spectral density (PSD) estimates were calculated on down-sampled data (1kHz). Theta power was calculated using 4000 sample Fourier transforms at a 0.25Hz resolution, while analysis gamma power was calculated using 500 sample Fourier transforms at 2.0Hz resolution, both using Hann windows with 50% overlap. Data were taken from channels within the expected range of impedance for a given probe type to allow exclusion of bad/broken channels (probe type 1: 1.1±0.3MΩ; probe type 2: 0.01-0.08MΩ (additional Z-coating)). The four channels closest to the optic fiber were excluded from all analyses, as they were most likely to be contaminated with opto-electric artefact [20]. Power spectra were then plotted for visual inspection blinded to recording condition. Recording blocks containing fewer than 1/4 of remaining channels (<7), after accounting for the removal of the four channels containing optogenetic artefacts from all datasets, were excluded from further analysis (n=4 sessions) as well as sessions with an abnormal power spectrum across all channels (n=5 sessions). Single recordings with spectral artefact were additionally excluded (n=1 recording from 1 session).

For frequency domain analyses, all power spectra computed over stimulation periods had the unstimulated baseline response subtracted to emphasise the effect of stimulation. The baseline spectrum for each channel was calculated by averaging unstimulated periods spread out throughout the session to account for drift. The baseline-subtracted spectra were first averaged across channels, then again along experimental variables (genotype, phase, amplitude) for visualisation. Subsequent, statistical comparisons focused on the mean values across specific frequency bands within the theta (4.5-7.5Hz) and gamma (56.25-93.75Hz) ranges.

## Statistical Analyses

Statistical analyses were undertaken using linear mixed effects (LME) models where possible (R: 4.3.2) to allow for appropriate analysis of longitudinal data with interactions and account for possible missing values. In the case there was insufficient data for a LME model, with the caveat that we were unable to control for repetition of subjects within and between sessions, paired t-tests were used as an alternative. While data was collected at two amplitudes (1 & 2mW), LME models demonstrated no significant differences between amplitudes in terms of main effect of amplitude or interaction terms at any given phase in either mouse group (ChR2; control), for each mouse type (PV-Cre; RBP4-Cre). Therefore, the 1 & 2mW data were combined into a ‘stimulation on’ condition. As a result, this means the same recording sessions contribute data in the analyses and plots that were collected for multiple amplitudes at each phase and stimulation pattern. Therefore, the number of recordings per phase may be greater than the number of sessions. In figure legends, the number of mice and recordings per phase are be provided.

Shapiro-Wilk normality tests demonstrated non-normality of the extracted PSD estimates, so data were z-scored prior to statistical analyses. Outliers within the theta range, identified as 1.5x the IQR for each subset of data were extracted and re-visually inspected. While there were no grounds for exclusion and all data was included for statistical analyses, these ‘outliers’ have been identified on plots accompanying statistics in a lighter colour for differentiation and were not included in plotting of boxplots. For all boxplots, grey boxes represent the interquartile range and median values from the data set, and the whiskers extend to minimum and maximum values calculated from their quartile ±1.5*IQR for each grouped session. Individual scatter points are session means for their respective group.

All LME models contained random effects of subject (each mouse) and date (denoting a session), to account for the variability within the data that arises from grouping of observations, as mice were used across several days and different mice were commonly recorded on the same date. This controls for variation in the random effects, while determining if there are associations between the fixed effects of the model. P-values were obtained by likelihood ratio tests and REML (restricted maximum likelihood) was set to false as ML (maximum likelihood) was used to estimate both fixed and random effects. To follow up significant interactions (p<0.05) pairwise t-tests were undertaken using the emmeans package in R (Tukey adjustment for multiple comparisons). Full results of LME models are given in supplementary tables 1-17.

## Results

To understand how phase-dependent stimulation modulates motor cortical theta and gamma oscillations, we developed a method for delivering optogenetic stimulation of the motor cortex triggered by specific phases of the ongoing theta oscillation in the motor cortical LFP in awake, head fixed mice. Stimulation was targeted to either PV-expressing inhibitory interneurons (PV-Cre) or Layer V excitatory pyramidal (RBP4-Cre) neurons (see Figs. S1 & S2). All statistical tests are LMEs (see methods) unless otherwise stated.

### Phase-dependent modulation of the motor cortical theta power by theta-phase targeted *continuous* optogenetic stimulation

We first evaluated the effect of theta-phase targeted, single continuous closed-loop optogenetic pulses (“continuous pulse”) on the power of the ongoing theta oscillation (Fig. 2). Optogenetic stimulation can be confounded by indirect effects of the applied light [21]. To ensure that effects were dependent on opsin-mediated activation of neuronal activity, we used control animals that had been injected with a construct that transfected Cre-dependent GFP, but not the opsin (GFP-controls). In opsin-transfected mice, we also compared real stimulation sessions to those when stimulation was turned off (i.e. phase is tracked, but stimulation is not applied and the timepoints at which stimulation *would* have been applied are use as the reference frame for analysis [STIM-ON and STIM-OFF]). We tested for significant main effects of group (ChR2 vs GFP-control), phase (peak, descending, ascending and trough) and stimulation (STIM-ON and STIM-OFF) and all interaction terms.

**Figure 2:**
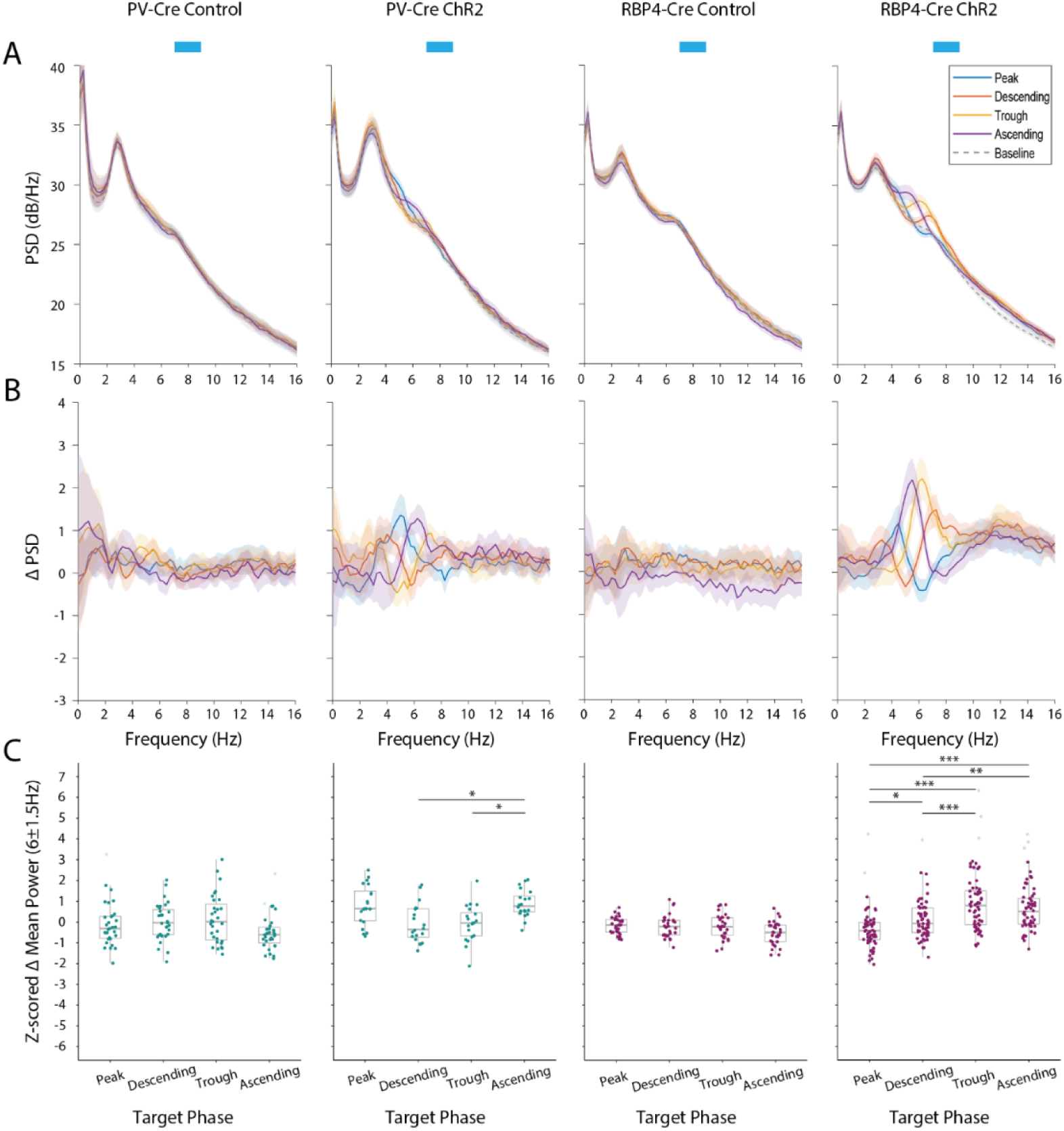
Closed-loop continuous stimulation of PV+ interneurons and RBP4+ pyramidal neurons led to phase-dependent effects on local theta power. (**A**) Mean power spectral density for PV-Cre & RBP4-Cre control (n=4 mice; 31 recordings/phase & n=4; 30 recordings/phase respectively) and ChR2 mice (n=5 mice; 20 recordings/phase & n=10; 60 recordings/phase respectively), targeting 4 different phases of the theta oscillation with continuous stimulation at 1 or 2mW. Shaded areas show ±2*SEM. (**B**) As in (A) but using baseline-subtracted spectra. (**C**) Boxplots showing the z-scored change in theta power with respect to baseline extracted between in the theta range (4.5-7.5Hz). Asterisks indicate significant differences in theta power between different target phases using LME, followed by pairwise post hoc t-tests with Tukey adjustment at p<0.05.

Stimulation of inhibitory interneurons using closed-loop continuous pulses led to significant phase-dependent modulation of theta power in ChR2 mice compared to GFP-controls (group*phase: F(3,288)=5.03, p=0.002; for full statistical analyses see Table S1; Fig. S3), and was only present under STIM-ON conditions (stimulation: F(1,288)=4.82, p=0.029). Given that we were primarily interested in stimulation-driven effects, we then tested phase dependence only in the STIM-ON condition (group, phase and their interaction - Table S2). This demonstrated significant phase-dependent modulation of theta power in ChR2 mice compared with GFP-controls (group*phase: F(3,184)=9.07, p<0.001). Post hoc-tests indicated significant differences in theta power between multiple phases in PV-Cre ChR2 mice, but not in the GFP-controls (Pairwise t-tests Tukey adjustment for multiple comparisons; Fig. 2C left).

In comparison, when stimulating excitatory RBP4+ neurons with the continuous protocol, there was a significant three-way interaction between group, phase and stimulation (F(3,469)=6.70, p<0.001; Table S3 & Fig. S3, right). Analysis of the STIM-ON data alone showed that ChR2 was necessary for phase-dependent theta power modulation (group*phase: F(3,323)=14.93, p<0.001; Table S4). Subsequent pairwise t-tests (Tukey adjustment for multiple comparisons) identified significant differences in theta power between multiple target phases in RBP4-Cre ChR2 mice, but not in the RBP4-Cre GFP-controls (Fig. 2C, right).

### Phase-dependent modulation of the motor cortical theta power with theta-phase targeted *gamma-frequency* optogenetic stimulation

Given the important role of theta-gamma coupling in the motor cortex [6,7,17], we hypothesised that gamma frequency optogenetic stimulation - a burst of three pulses at 75Hz (gamma pulse) - might be more effective in modulating theta power than the continuous pulse (Fig. 3). In PV-Cre mice, group and phase had significant effects on theta power (group*phase: F(3,324)=4.29, p=0.006) during closed-loop gamma pulse stimulation (Table S5 & Fig. S4). For STIM-ON data, we identified the same significant interaction (group*phase: F(3,205)=6.31, p<0.001; Table S6) and post hoc tests confirmed significant differences in theta power between multiple phases in PV-Cre ChR2 mice (pairwise t-tests Tukey adjustment, Fig. 3C, left).

**Figure 3:**
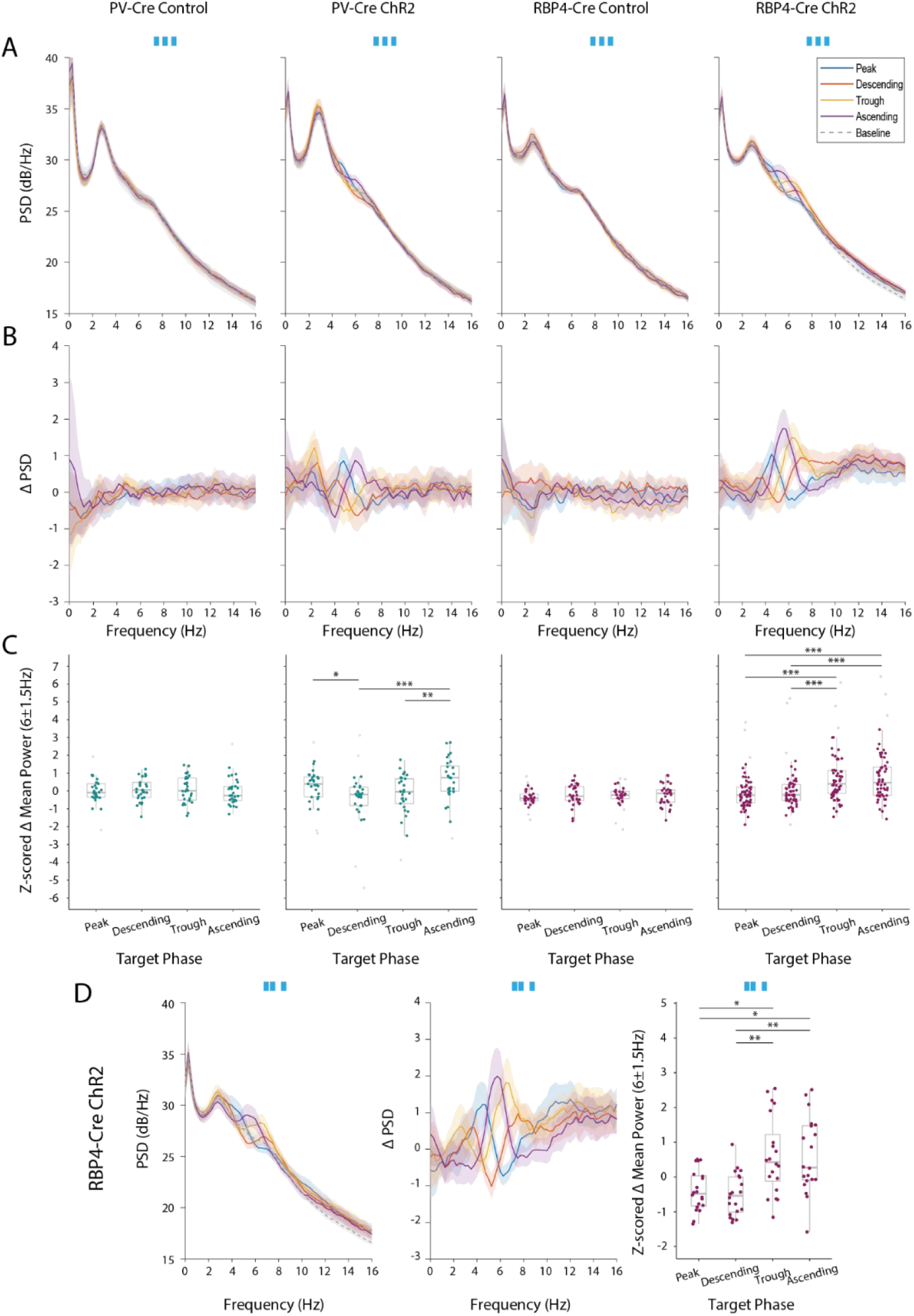
**Closed-loop gamma stimulation of PV+ interneurons and RBP4+ pyramidal neurons lead to phase-dependent effects on local theta power**. (**A**) Mean power spectral density for PV-Cre & RBP4-Cre GFP (n=4 mice; 29-30 recordings/phase & n=4; 30 recordings/phase respectively) and ChR2 mice (n=7 mice; 28-29 recordings/phase & n=10; 60 recordings/phase respectively) targeting 4 different phases of the theta oscillation with gamma stimulation at 1 or 2mW. Shaded areas show ±2*SEM. (**B**). As in (A) but using baseline-subtracted spectra. (**C**) Boxplots with the z-scored change in theta power with respect to baseline extracted between in the theta range (4.5-7.5Hz). Asterisks indicate significant differences in theta power between different target phases using LME, followed by pairwise post hoc t-tests with Tukey adjustment at p<0.05. (**D**) As in A-C, but for RBP4 ChR2 mice (n=5; 20 recordings/phase) receiving arrhythmic gamma stimulation.

Gamma pulse stimulation of excitatory neurons also led to significant phase-dependent modulation of theta power dependent on group and phase (group*phase: F(3,554)=2.65, p=0.048). As might be expected, theta modulation varied with stimulation condition, but only in the RBP4+ ChR2 (group*stimulation: F(1,545)=25.45, p<0.001; Table S7 & Fig. S4). We therefore performed an LME model for just the STIM-ON data (Table S8). A significant group*phase interaction (F(3,324)=5.82, p<0.001) demonstrated the necessity of ChR2 for phase-dependent theta modulation. Subsequent pairwise t-tests (Tukey adjustment for multiple comparisons) identified significant differences between theta power at multiple target phases (Fig. 3C, right).

To determine the importance of rhythmicity of gamma pulse stimulation on modulation of theta power, in RBP4-Cre ChR2 mice we tested a control stimulation pulse without the oscillatory component of the stimulus (“arrhythmic gamma”). For this control protocol, the timing of the middle of the three pulses was pseudorandomly placed between the two outside pulses. Thus, the total number of pulses and mean intervals between pulses were identical to gamma pulse stimulation, but without gamma rhythmicity. We tested the effects of stimulation pattern (gamma pulse, arrhythmic gamma), stimulation (on, off), phase (peak, descending, trough & ascending) and all combinations of interactions (Table S9). There were no effects of stimulation pattern on phase-dependent theta power modulation (stimulation pattern*phase: F(3,299)=0.61, p=0.609), suggesting that the rhythmic component of the pulses was not necessary to modulate theta power. Supporting this finding, a comparison of all three optogenetic stimulation patterns in the STIM-ON data revealed no significant interaction terms (Table S10). Follow up pairwise t-tests (Tukey adjustment for multiple comparisons) identified significant differences between theta power at multiple phases following arrhythmic gamma frequency stimulation (Fig. 3D). Overall, these results show that while bursts of optogenetic pulses applied over a specific phase of the theta cycle were sufficient for phase dependent modulation of theta power, a periodic/stable gamma frequency was not necessary to achieve these effects.

### Closed-loop interaction is necessary for phase-dependent effects on theta power

Altering the target phase of closed-loop stimulation can produce different rates and patterns of stimulation pulses, due to the continuous interaction between the neural signal and stimulus output [8]. These primary statistical features of the stimulation trains could result in different effects between target phases. As we have done previously, to ensure phase-dependent theta modulation resulted from closed-loop interaction with the underlying oscillation, we used an open-loop replay paradigm whereby the stimulation patterns generated by closed-loop stimulation at each target phase were “played back” in the same session they were recorded [8,9] (Fig. 1E). Open-loop replay stimulation did not reproduce phase-specific effects seen in closed-loop recordings in RBP4-Cre or PV-Cre ChR2 mice (Fig. 4 & S5). We compared the z-scored difference in mean theta power between the most and least amplifying phases during closed-loop stimulation and matched open-loop counterparts in RBP4-Cre ChR2 mice (LME: Stimulation type; gamma frequency: F(1,46)=61.68, p<0.001, Fig. 4D left; continuous pulse: F(1, 25)=34.16, p<0.001, Fig. 4D, right). Where the datasets for these recordings in RBP4-Cre and PV-Cre ChR2 mice were too small for LME models, the same data were compared using paired t-tests following arrhythmic gamma (t(8)=4.42, p=0.01 (Fig. S5F)) and continuous stimulation (t(4)=4.55, p=0.01 (Fig. S5E)), respectively.

**Figure 4:**
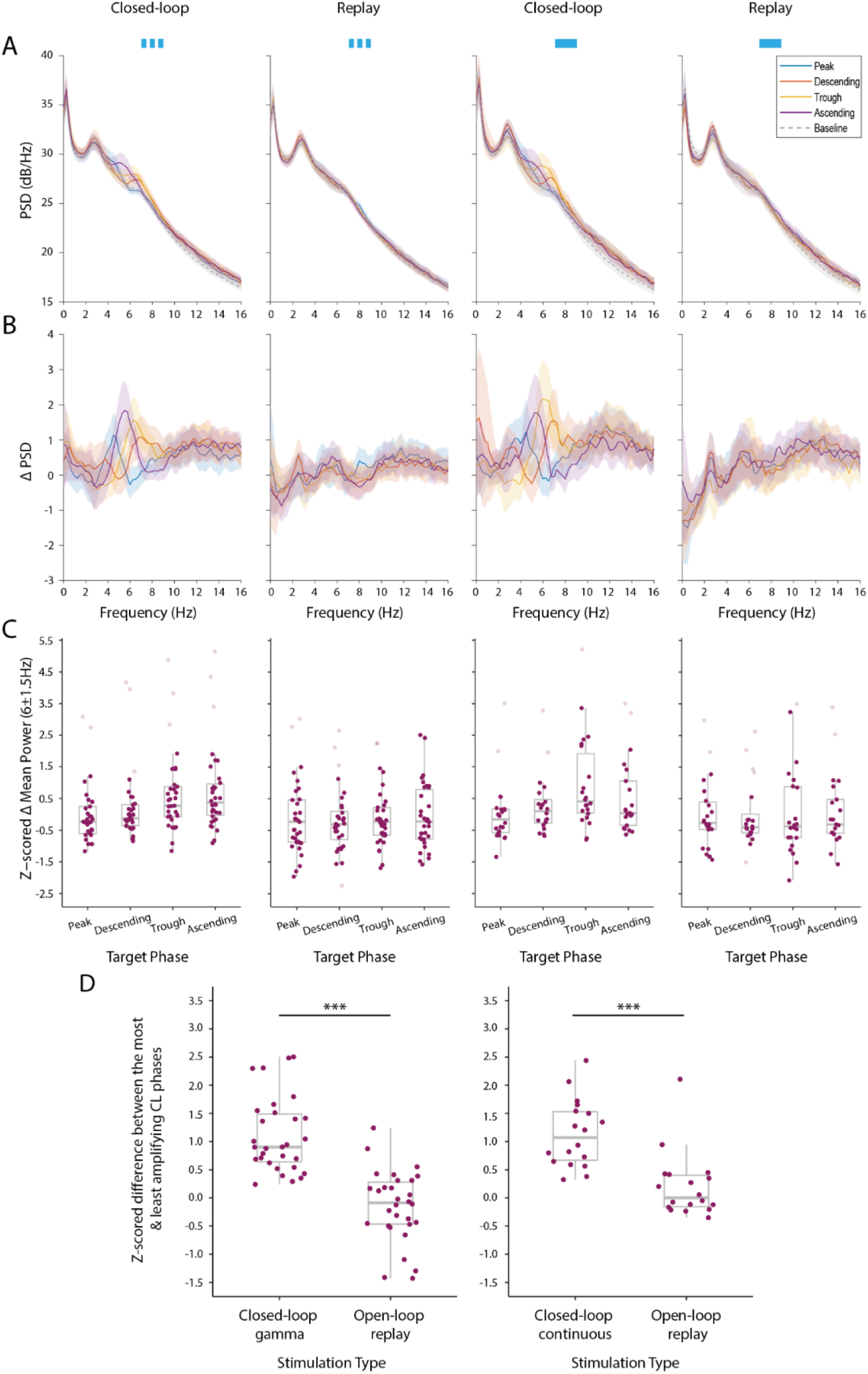
Closed-loop interaction is necessary in producing the same theta phase-dependent effects following both gamma and continuous optogenetic stimulation. **(A)** Mean power spectral density in RBP4-Cre ChR2 mice with gamma (n=9 mice; 32-34 recordings/phase) or continuous (n=4 mice; 20-22 recordings/phase) at 1 or 2mW using closed-loop and open-loop replay (replay) protocols. Shaded areas show ±2*SEM. **(B)** As in (A) but using baseline-subtracted spectra. (**C**) Boxplots with the z-scored change in theta power with respect to baseline extracted between in the theta range (4.5-7.5Hz). **(D)** For each main stimulation pattern (gamma - left/continuous - right) the difference between the most and least amplifying phase for a closed-loop recording are plotted as individual scatter points, the difference between the same phases is then taken in the open-loop recording and displayed as individual scatter points. Significant differences (*p<0.05) were found between the z-scored difference in theta power of the most and least amplifying phases in each specific closed-loop recording and their open-loop counterpart for both gamma and continuous stimulation (LME, main effect stimulation type). In these specific statistical analyses only sessions with a complete data set for all four phases at the same amplitude were used.

### Closed-loop stimulation of cortical pyramidal neurons leads to greater amplification of gamma power than PV-expressing interneurons

Theta and gamma oscillations are often coupled in cortical circuits and gamma frequency stimulation has previously been shown to modulate gamma power [17,22–25]. We next investigated whether our stimulation protocols also modulated gamma-frequency oscillations (Fig. 5; S6-9). Mean power spectra in the gamma range for different target phases using both gamma (Fig. 6 & Fig. S6-7) and continuous (Fig. S8-9) stimulation were tightly overlayed, suggesting little difference in gamma modulation due to theta phase targeting. In line with this observation, we found no significant effects of target theta phase in any of the stimulation protocols (Tables S11-14).

**Figure 5:**
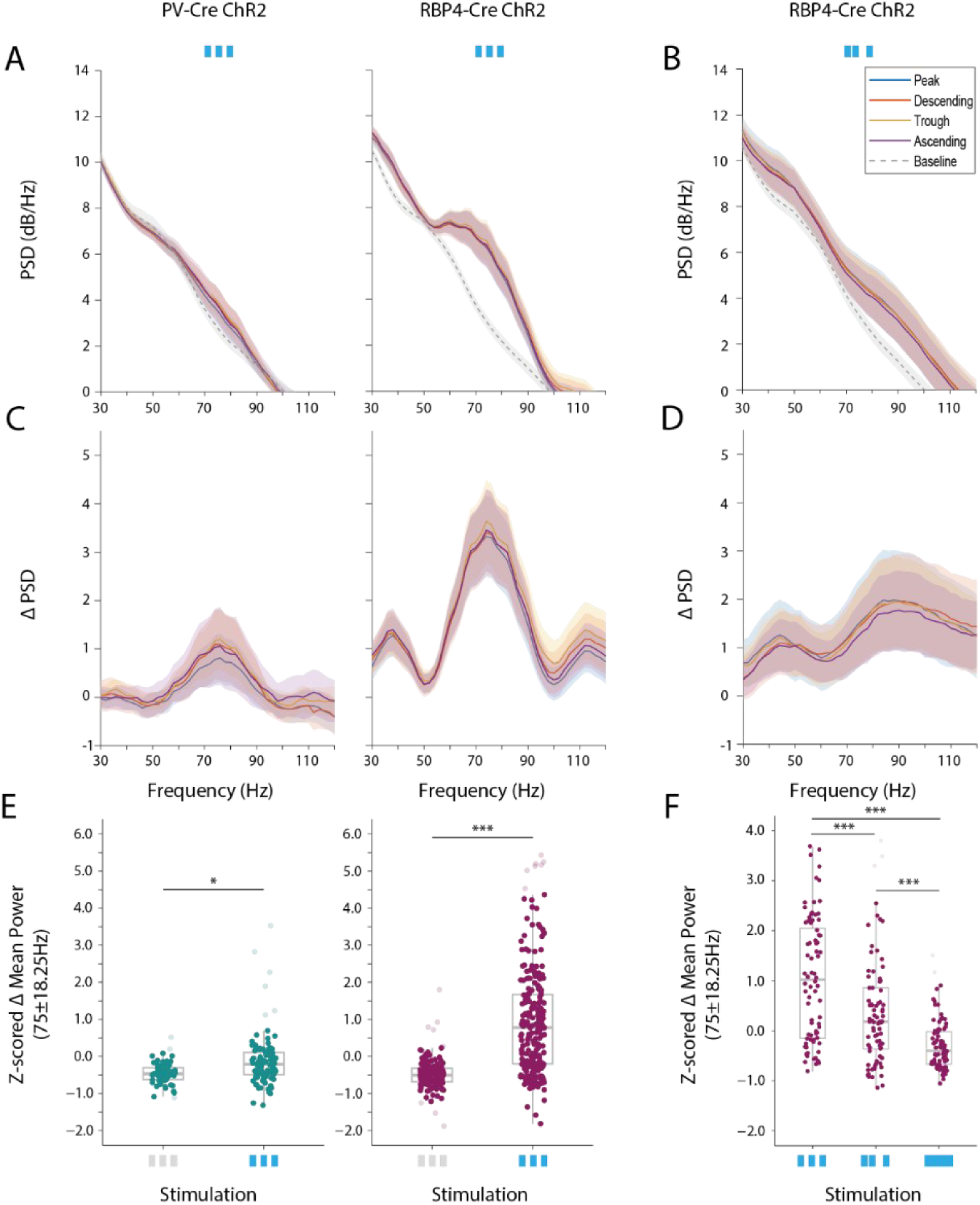
**Closed-loop gamma stimulation of PV+ interneurons and RBP4+ pyramidal neurons led to phase-dependent effects on local gamma power**. **(A)** Mean power spectral density from 30-120Hz for PV-Cre ChR2 (n=6 mice; 26-27 recordings/phase) and RBP4-Cre ChR2 (n=10; 60 recordings/phase) mice, targeting four different phases of the theta oscillation with gamma stimulation at 1 or 2 mW. Shaded areas show ±2*SEM. **(B)** Mean power spectral density from 30-120Hz for RBP4-Cre ChR2 mice receiving arrhythmic gamma stimulation (n=5; 20 recordings/phase) at 2mW. **(C-D)** As in (A-B) but using baseline-subtracted spectra. **(E)** Statistical analyses performed demonstrated no significant main effect or interaction terms including phase on gamma power (all p>0.05; LME), subsequently, data from all phases was combined for further analyses. Boxplots with the z-scored change in mean gamma power with respect to baseline extracted between in the gamma band (56.75-93.25Hz) for PV-Cre and RBP4-Cre ChR2 mice during gamma stimulation (1 or 2mW) vs no stimulation. Gamma power was significantly higher when stimulation was on in both PV-Cre and RBP4-Cre mice (*p=0.048 & p<0.001 respectively, LME – significant group*stimulation interaction; followed by pairwise post hoc t-tests with Tukey adjustment). **(F)**. Z-scored change in mean gamma power in RBP4-Cre mice was significantly different between all three stimulation patterns (2mW only; *p<0.001 LME followed by pairwise post hoc t-tests with Tukey adjustment).

**Figure 6:**
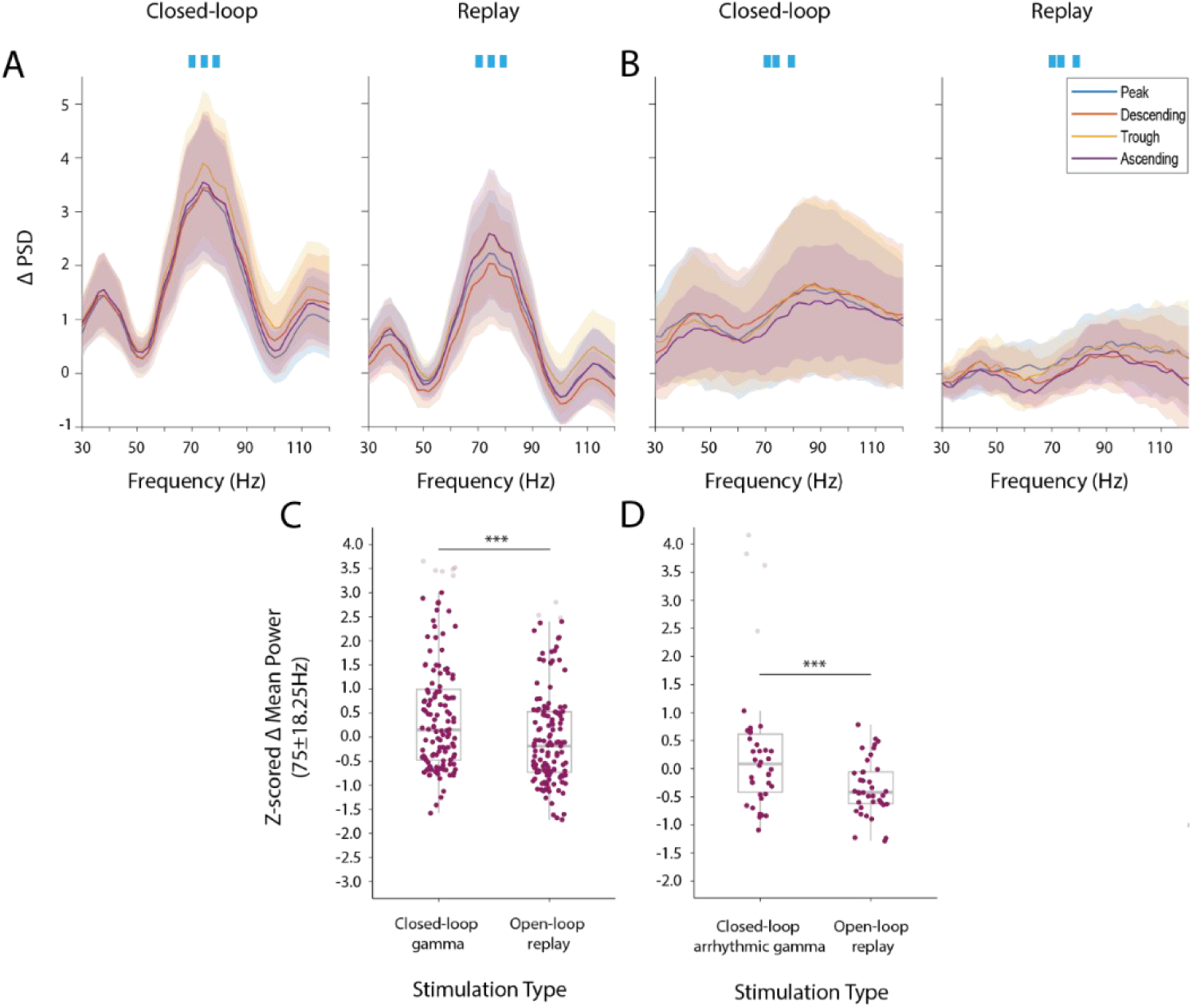
**Closed-loop interaction increases gamma amplification in RBP4-Cre mice**. (**A**) Baseline-subtracted mean power spectral density from 30-120Hz in RBP4-Cre ChR2 mice receiving closed-loop gamma (n=9 mice; 32-34 recordings/phase, 132 total recordings; 1 or 2mw) or arrhythmic gamma (n=5; 9-10 recordings/phase, 38 total recordings; 2mW only) using both closed-loop and open loop replay protocols. **(C-D)** Boxplots demonstrate z-scored change in gamma power with respect to baseline extracted between our target range of 56.75-93.25Hz (75Hz±18.25). Gamma power was significantly higher during closed-loop protocols for both gamma and arhythmic gamma stimulation (p<0.001 (LME model: group (ChR2, control)*stimulation (on, off)) followed by pairwise post hoc t-tests with Tukey adjustment).

As there was no phase-dependence, we combined data across target phases to examine the effects of different stimulation protocols on gamma power. We first tested for significant effects of group (PV-Cre (ChR2, GFP-control) and RBP4-Cre mice (ChR2, GFP-control)) and stimulation (on, off), alongside their interaction term for each stimulation pattern (gamma (Table S15); continuous (Table S16)). Both gamma (group*stimulation: F(3,861)=50.14, p<0.001; Table S15, Fig. 5 & S6-7) and continuous pulse (group*stimulation: F(3,751)=7.93, p<0.001; Table S16, Fig. S8-9) stimulation protocols modulated gamma power, although continuous stimulation did not lead to the same clear peak in the gamma range (Fig. S8). Post hoc tests confirmed a significant difference between gamma pulse stimulation being on and off in the ChR2 group (Fig. 5 & S6), for both PV-Cre (t(865.2)=-3.051, p=0.048) and RBP4-Cre mice (t(866.3)=-20.385, p<0.001) (Fig. 5E). However, while there was a significant effect of gamma stimulation in the STIM-ON condition between RBP4-Cre ChR2 and GFP-controls (t(29.4)=-4.675, p=0.001), this was not the case for PV-Cre ChR2/GFP-controls (t(39.3)=-0.322, p>0.05) (Fig. 5 & S7). This suggests that any modulation of gamma in PV-Cre animals was not stimulation/opsin mediated. Post hoc tests only identified a significant difference in gamma power following continuous stimulation (on versus off) in RBP4-Cre ChR2 mice (t(785.9)=-7.145, p<0.001; Fig. S8C). Overall, gamma stimulation of excitatory neurons was the most effective way to drive gamma power in the LFP. Moreover, in RBP4-Cre ChR2 mice, arrhythmic gamma stimulation was significantly less effective in modulating gamma power (LME model: stimulation pattern (gamma, arrhythmic gamma & continuous) - F(2,227)=62.75, p<0.001). Follow up post hoc tests identified significant differences between each combination of stimulation patterns (all p<0.001, Fig. 5F). While the rhythmicity of the gamma pulse did not affect the modulation of theta power, it did therefore affect the amount of gamma amplification.

### Theta-phase locking increases amplification of gamma power, but not in a phase-dependent manner

We hypothesised that if theta-phase targeting has no effect on gamma amplification, open-loop replay should have equivalent effects to closed-loop stimulation on this parameter. Using the gamma pulse protocol in RBP4-Cre mice, we found that this was not the case, rather that gamma power modulation was stronger for closed-loop stimulation than open-loop replay of the same patterns (Fig. 6). This was revealed by a significant effect of group (ChR2, control) by stimulation type (closed-loop, open-loop replay) interaction (F(1,366)=14.03, p<0.001; Table S17) followed by pairwise post hoc t-tests with Tukey adjustment (Fig. 6C). A similar result was found for arrhythmic gamma stimulation (main effect of stimulation type LME; F(1,64)=23.59, p<0.001, Fig. 6D) in ChR2 mice. In summary, both closed-loop phase targeting and the rhythmicity of the gamma pulse influenced amplification of gamma power by our stimulation protocols.

## Discussion

Here we demonstrate that closed-loop, theta phase-targeted optogenetic stimulation of inhibitory interneurons and excitatory pyramidal neurons in the mouse motor cortex results in phase-dependent enhancement of theta power. Gamma-pulse stimulation was sufficient to modulate theta power, despite considerably less energy being applied during the target phase of stimulation. Closed-loop interaction was necessary for producing theta modulation, which was not reproduced by replaying the same stimulation patterns uncoupled from ongoing activity. Stimulating pyramidal neurons using gamma stimulation led to the largest amplification of gamma power. While the amplification of gamma modulation was not theta-phase dependent, it was larger during phase-targeted/closed-loop stimulation *per se*. Overall, we demonstrate the feasibility and effectiveness of this method to explore the mechanisms underlying the response of cortical microcircuits to closed-loop stimulation paradigms.

### Theta power is amplified at different target phases for stimulation of pyramidal neurons and interneurons

Hitherto, studies into phase-targeted stimulation have either used electrical stimulation or have stimulated a single neuron type with optogenetics. Neither of these approaches gives insight as to how different components of the underlying neural circuits respond to the same closed-loop approach. Our work reveals an offset of 90° between stimulation phases that amplify the theta oscillation between our two target neuronal populations. Maximal amplification of theta occurred for stimulation of interneurons and excitatory neurons at the ascending/peak and trough/ascending LFP phases, respectively. This finding is in line with predictions from computational models exploring perturbations of excitatory and inhibitory circuits [26]. Moreover, studies targeting gamma oscillations within brain regions such as the hippocampus and prefrontal cortex have identified that the peak firing phase of pyramidal neurons precedes that of interneurons [27–30]. Identifying how individual neurons in these different populations respond to closed-loop stimulation will provide further insight as to how the microcircuit responds to these approaches.

Alongside phase-dependent amplification, frequency shifts in theta were observed when targeting both inhibitory and excitatory cell types. This may result from a constant phase-reset of the theta oscillation, in which an oscillation alters its timing to synchronise with the external disrupting stimulus [31,32]. Ultimately, whether optogenetic stimulation advances or delays the theta oscillation appears to depend on the phase targeted. The magnitude of a phase-reset has additionally been found to depend on oscillatory amplitude, as well as input strength, in which lower oscillatory power can lead to larger phase shifts [32–34]. This aligns with the larger frequency shift (∼1Hz) observed when targeting interneurons where the amplitude is less modulated, in comparison to pyramidal neurons where a smaller frequency shift is demonstrated (∼0.5Hz). In contrast to our previous studies [8,9], we were unable to bidirectionally modulate theta in this study. This could in part be explained by the low baseline amplitude of the motor cortical theta power, leaving only a small lower dynamic range in which to achieve suppression.

### Stimulation of pyramidal neurons led to greater amplification of gamma power

Contrary to previous studies highlighting a role of parvalbumin containing interneurons in the generation and maintenance of gamma oscillations [22,23,35], stimulation of excitatory neurons was more effective in amplifying gamma power. More specifically, rhythmic gamma stimulation (75Hz) of excitatory, but not inhibitory, neurons led to significant enhancement of gamma oscillations compared to controls. This is supported by Nicholson and colleagues (2018) [12] who demonstrate bidirectional modulation of gamma oscillations through optogenetic stimulation of pyramidal cells, but not interneurons, *in vitro.* They proposed that this resulted from out-of-phase recruitment of interneurons powerfully suppressing the oscillation [12], which could be a direct result of desynchronisation. This mechanism could potentially explain our findings, where stimulation induced amplification of gamma power is masked by circuit interactions. Our study has the potential to add to this line of research as further analyses of unit data could directly determine the extent of gamma oscillatory phase locking for individual cell types. In addition, Sohal and colleagues (2009) found optogenetic stimulation of pyramidal neurons evoked gamma oscillations that are strongly phase-locked to the stimulation [23]. Overall, this finding suggests that cortical gamma power may be more effectively driven through stimulation of excitatory neurons.

### Concluding remarks

In summary, our work demonstrates the potential of phase-locked optogenetic stimulation to precisely modulate microcircuit-level mechanisms underlying specific oscillatory dynamics. This approach can be utilised to inform development of brain stimulation methods through underlying processes that modulate these activities under pathological conditions in humans.

### Limitations of the study

While head-fixed preparations enable precise control over recording and stimulation conditions, by definition they reduce the capacity for movement. Given that our recordings were in motor cortex, responses to stimulation could differ under conditions where spontaneous and/or goal-directed movements were present. In human studies, increased theta activity was related to suppression of movement [36,37], which may align with the presence of theta when movement is suppressed in a head-fixed condition in mice. However, increases in gamma power have been observed during motor preparation and movement [38–40] and may be lower under these conditions. Therefore, this study does not fully capture the dynamic modulation or complexity of these oscillations under well-defined movement conditions. Optogenetic stimulation can induce photoelectrical artefacts, tissue heating effects, and non-specific neuronal responses that may lead to unintended false positive results [21,41–43]. We have carefully controlled for such effects using control mice and no stimulation conditions, using low laser powers and by visually inspecting the data to ensure that our reported results are genuinely observed effects. Nevertheless, we cannot completely rule out the influence of such artifacts in our LFP recordings.

Finally, while we were successful in using closed-loop optogenetics to manipulate the motor cortical theta oscillation, there are obvious constraints in translating these methods to humans. However, closed-loop optogenetics can play a key role in informing and advancing future therapeutics, elucidating the role of different neurons in the microcircuitry.

## Supporting information

Supplemental-Statistical-Tables

