## Supplemental-Figures for "Modulation of motor cortical theta and gamma oscillations using phase-targeted, closed-loop optogenetic stimulation of local excitatory and inhibitory neurons"

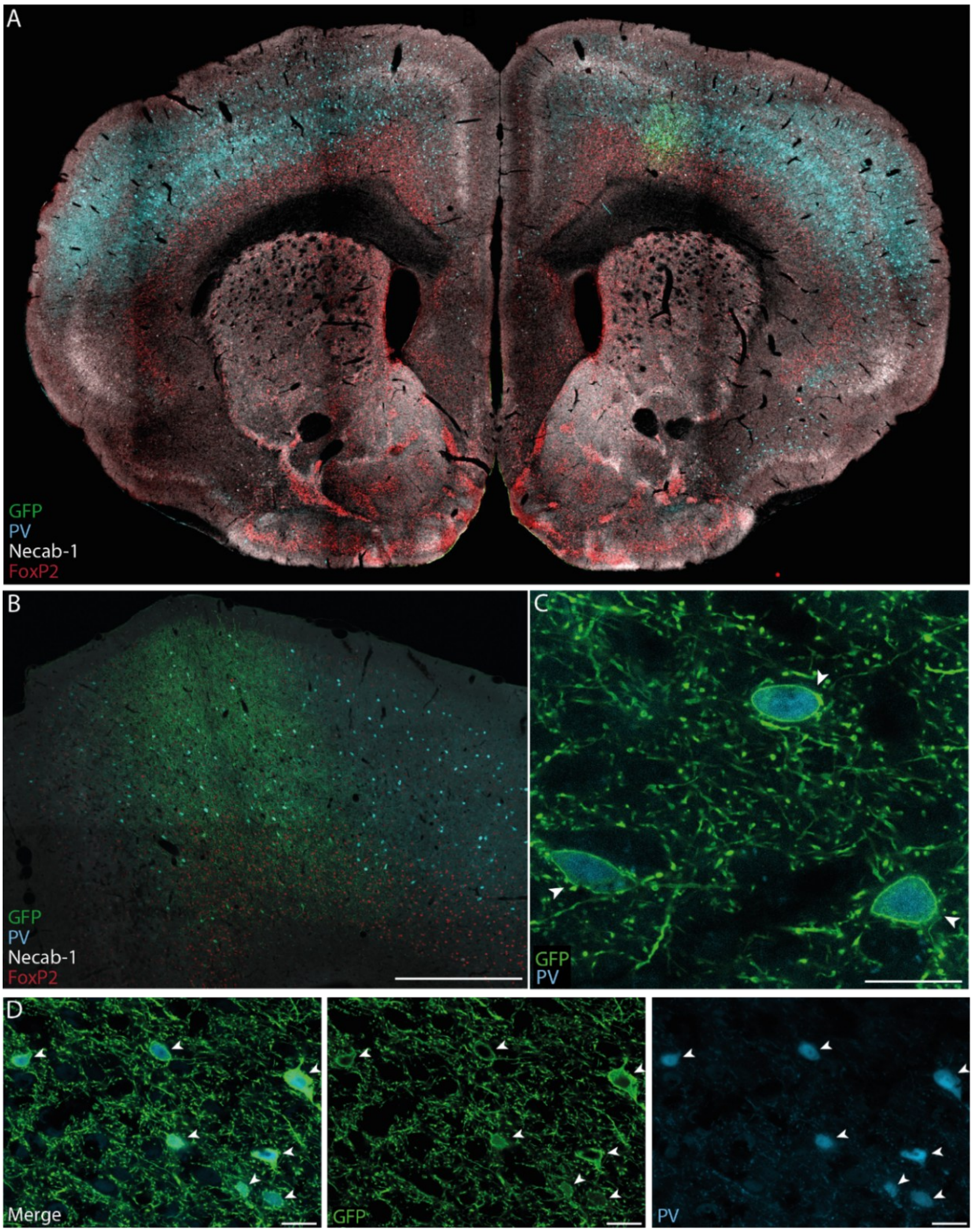


**S1: ChR2-GFP expression was localised to PV-expressing neurons in PV-Cre mice.** PV-Cre mice were injected either with a Cre-dependent control virus (GFP) or a virus containing ChR2-GFP. Low magnification (5x) image of PV, ChR2-GFP, Necab-1 and FoxP2 immunoreactivity on a whole brain coronal section (**A**) & within the cortex (**B**; 20x magnification, scale bar = 500mm). The Cre-dependent virus remains within the cortex and does not spread to deeper structures. (**C** & **D**) Higher magnification confocal images of ChR2-GFP & PV expression from a section of layer 5 within the motor cortex. Scale bar represents 20mm. There is extensive overlap between cells expressing both green fluorescent protein (GFP) and the calcium-binding protein Parvalbumin (PV), identified by white arrows. This confirms expression of PV-Interneurons with the Cre-dependent ChR2 virus.

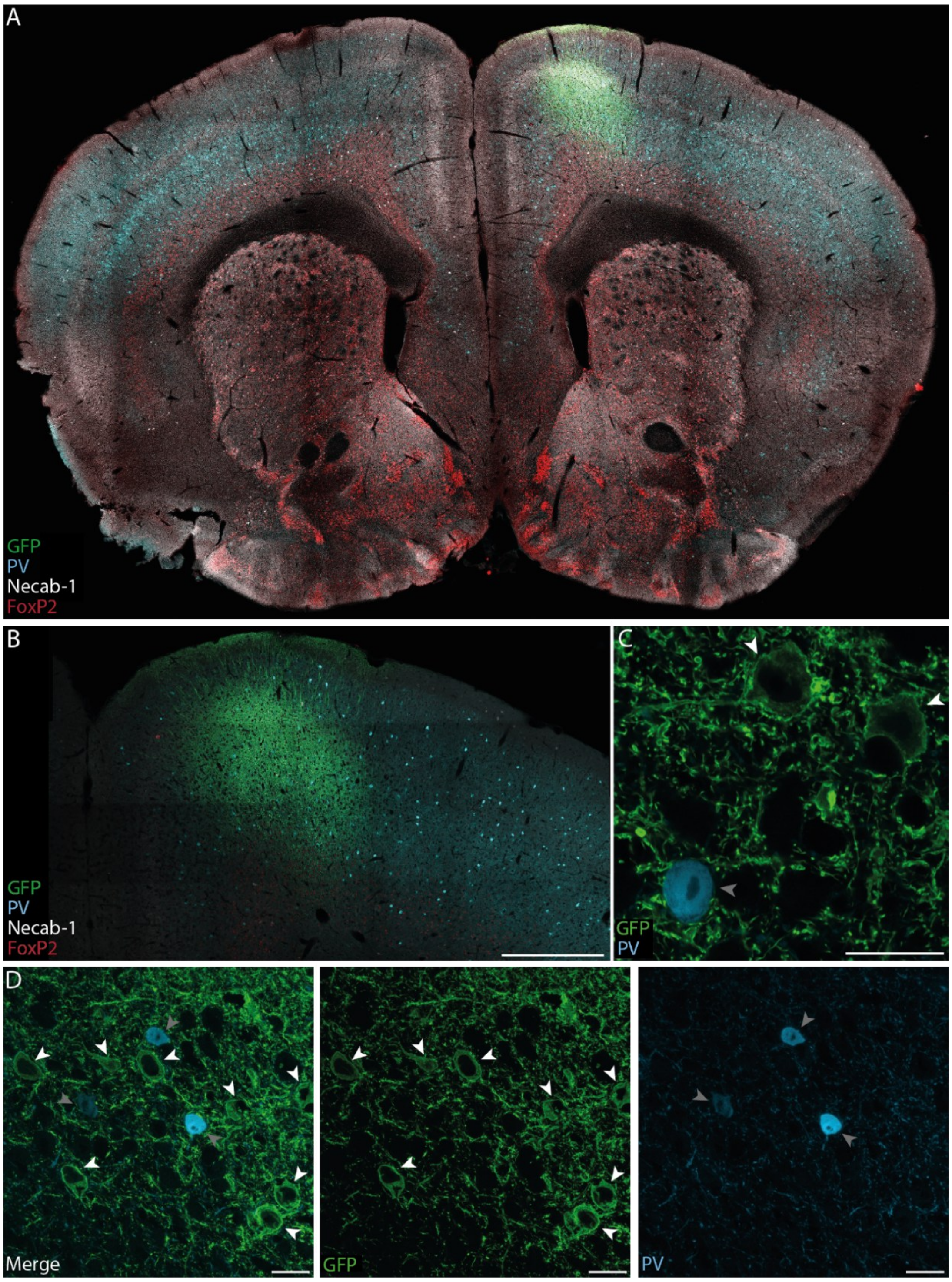


**S2: Localised expression of ChR2-GFP and no overlap with PV-expressing neurons in RBP4-Cre mice.** RBP4-Cre mice were injected either with a Cre-dependent control virus (GFP) or a virus containing ChR2-GFP. Low magnification (5x) image of PV, ChR2-GFP, Necab-1 and FoxP2 immunoreactivity on a whole brain coronal section (**A**) & within the cortex (B; 20x magnification, scale bar = 500mm). The Cre-dependent virus remains within the cortex and does not spread to deeper structures, although demonstrates connections to the contralateral hemisphere (**B**). Higher magnification images (63x) of ChR2-GFP & PV expression from a section of layer 5 within the motor cortex (**C & D**; scale bar = 20mm). There is no neurochemical marker to identify the presence of RBP4, instead, we demonstrate separation between cells identified as GFP expressing pyramidal neurons expressing and inhibitory interneurons expressing the calcium-binding protein Parvalbumin (PV) - Arrows: white – GFP; grey – PV.

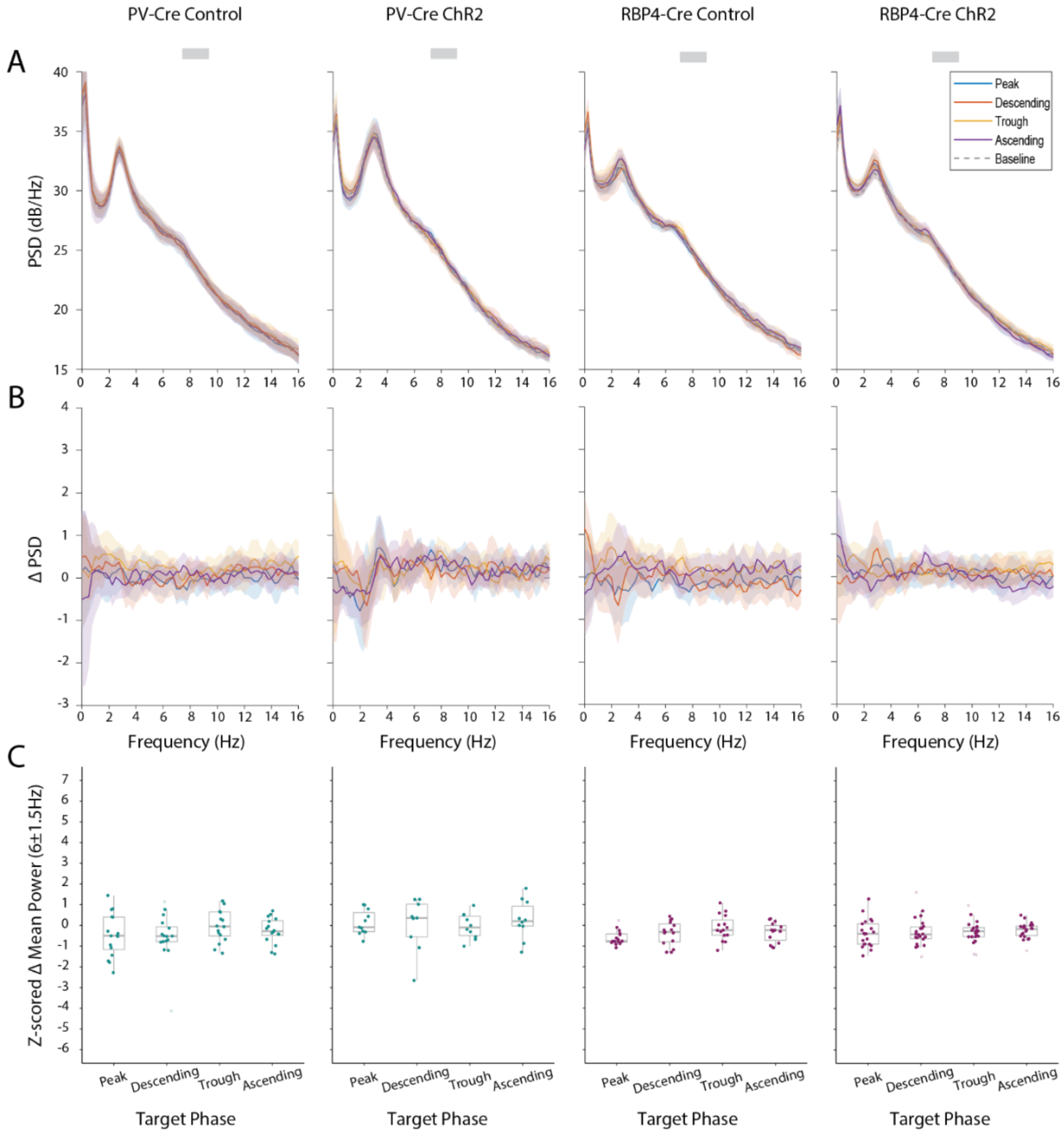


**S3: Closed-loop unstimulated continuous of PV+ interneurons and RBP4+ pyramidal neurons led to no phase-dependent effects on local theta power.** (**A**) Mean power spectral density for PV-Cre & RBP4-Cre GFP-control (n=4 mice; 16 recordings/phase & n=4; 15 recordings/phase) and ChR2 mice (n=5 mice; 9-11 recordings/phase & n=6; 21-23 recordings/phase), targeting 4 different phases of the theta oscillation with continuous stimulation when stimulation was turned off (0mW) (i.e. using reference time points when stimulation would have been applied). Shaded areas show ±2*SEM. (**B**). As in (A) but using baseline-subtracted spectra (**C**). Boxplots showing the z-scored change in theta power with respect to baseline extracted between in the theta range (4.5-7.5Hz). No significant differences (p>0.05) in theta power were identified between different target phases when unstimulated using LME – PV-Cre: significant group*stimulation interaction & RBP4-Cre: significant group*phase*stimulation interaction; followed by pairwise post hoc t-tests with Tukey adjustment.

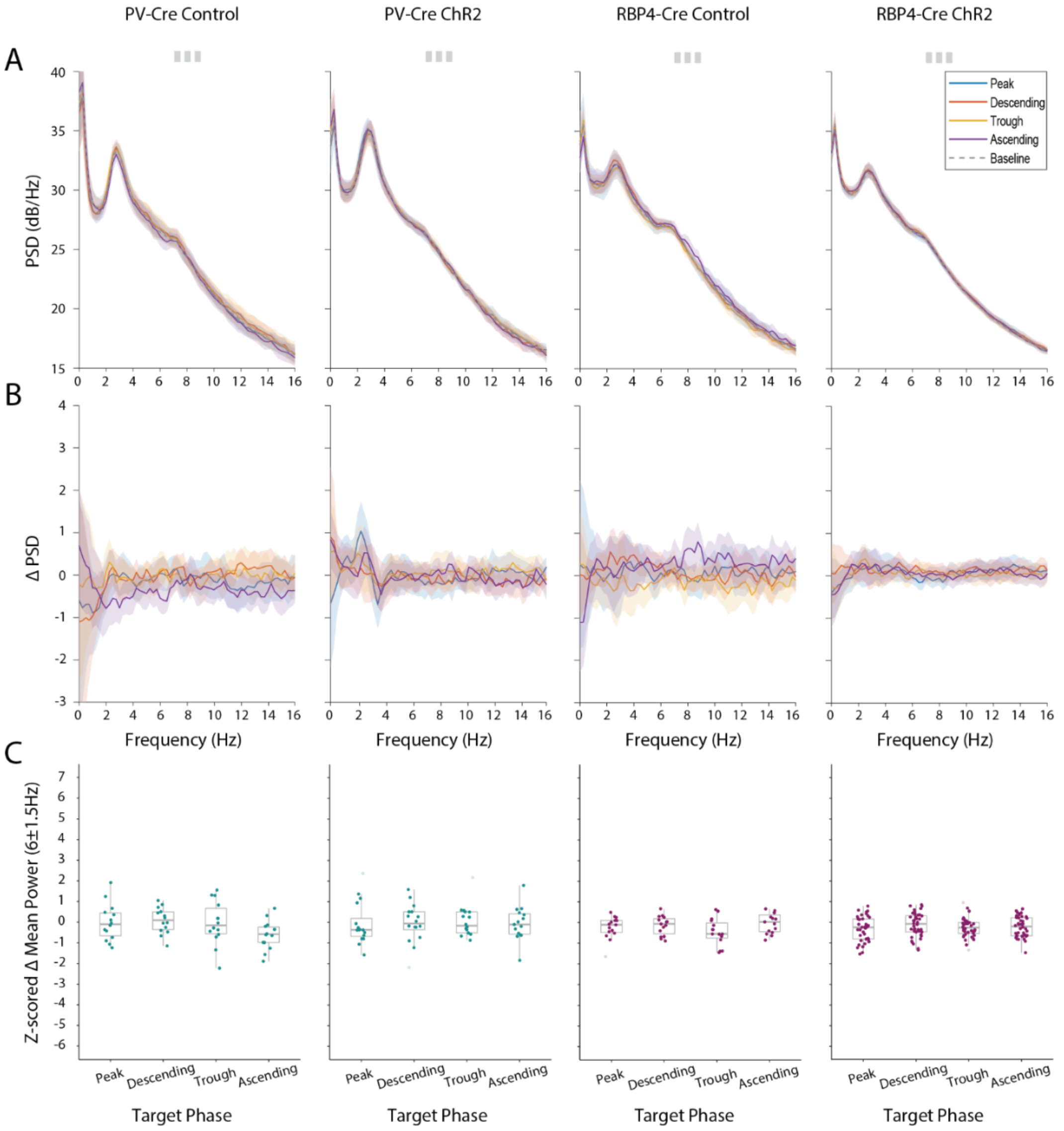


**S4: Closed-loop unstimulated gamma of PV+ interneurons and RBP4+ pyramidal neurons led to no phase-dependent effects on local theta power.** (**A**) Mean power spectral density for PV-Cre & RBP4-Cre GFP-control (n=4 mice; 14 recordings/phase & n=4; 15 recordings/phase) and ChR2 mice (n=7 mice; 15-16 recordings/phase & n=10; 40 recordings/phase), targeting 4 different phases of the theta oscillation with gamma stimulation when stimulation was turned off (0mW) (i.e. using reference time points when stimulation would have been applied). Shaded areas show ±2*SEM. (**B**). As in (A) but using baseline-subtracted spectra (**C**). Boxplots showing the z-scored change in theta power with respect to baseline extracted between in the theta range (4.5-7.5Hz). No significant differences ($p>0.05$) in theta power were identified between different target phases when unstimulated using LME – significant group*stimulation interaction; followed by pairwise post hoc t-tests with Tukey adjustment in PV-Cre or RBP4-Cre mice.

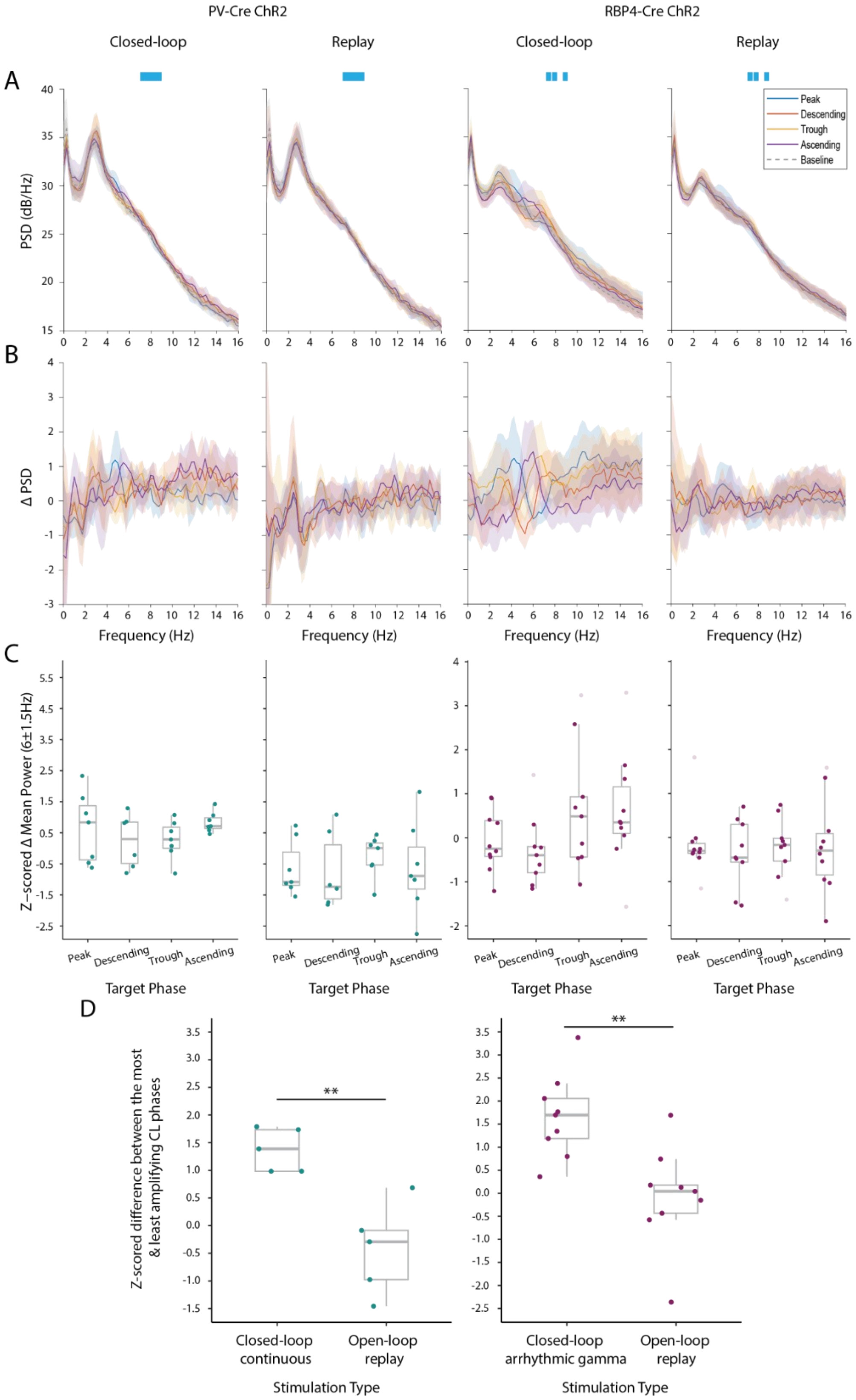


**S5: Closed-loop interaction is necessary in producing the same theta phase-dependent effects following continuous optogenetic stimulation of PV+ interneurons and arrhythmic gamma stimulation of RBP4+ pyramidal neurons. (A)** Mean power spectral density in PV-Cre ChR2 with continuous (n=3 mice; 6-7 recordings per phase) and RBP4-Cre ChR2 mice with arrhythmic gamma (n=5 mice; 9-10 recordings/phase) at 1 or 2mW using closed-loop and open-loop replay (replay) protocols. Shaded areas show ±2*SEM. **(B)** As in (A) but using baseline-subtracted spectra. (**C**)

Boxplots with the z-scored change in theta power with respect to baseline extracted between in the theta range (4.5-7.5Hz). **(D)** For each main stimulation pattern (continuous - left/arrhythmic gamma - right) the difference between the most and least amplifying phase for a closed-loop recording are plotted as individual scatter points, the difference between the same phases is then taken in the open-loop recording and displayed as individual scatter points. Significant differences (*$p<0.05$) were found between the z-scored difference in theta power of the most and least amplifying phases in each specific closed-loop recording and their open-loop counterpart for both continuous and arrhythmic gamma stimulation (paired t-test). In these specific statistical analyses only sessions with a complete data set for all four phases at the same amplitude were used.

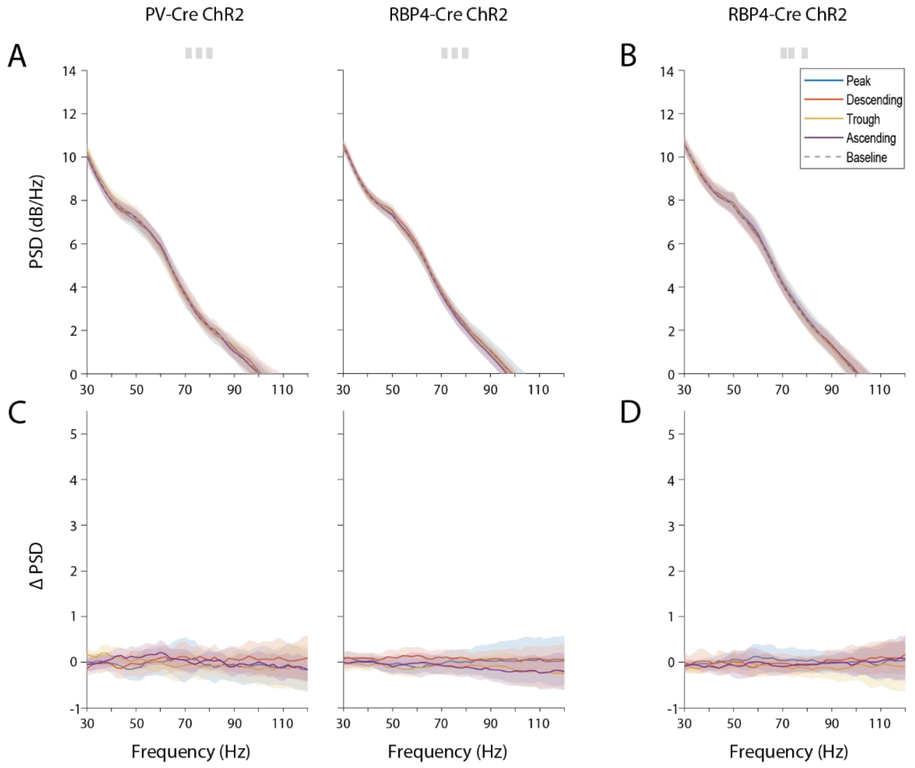


**S6: Closed-loop unstimulated gamma of PV+ interneurons and RBP4+ pyramidal neurons led to no phase-dependent effects on local gamma power. (A)** Mean power spectral density from 30-120Hz for PV-Cre ChR2 (n=6; 14-15 recordings/phase) and RBP4-Cre ChR2 (n=10: 40 recordings/phase) mice, targeting four different phases of the theta oscillation with gamma stimulation when stimulation was turned off (0mW) (i.e. using reference time points when stimulation would have been applied). Shaded areas show ±2*SEM. **(B)** Mean power spectral density from 30-120Hz for RBP4-Cre ChR2 mice receiving arrhythmic gamma stimulation (n=5; 19 recordings/phase) at 0mW. **(C-D)** As in (A-B) but using baseline-subtracted spectra.

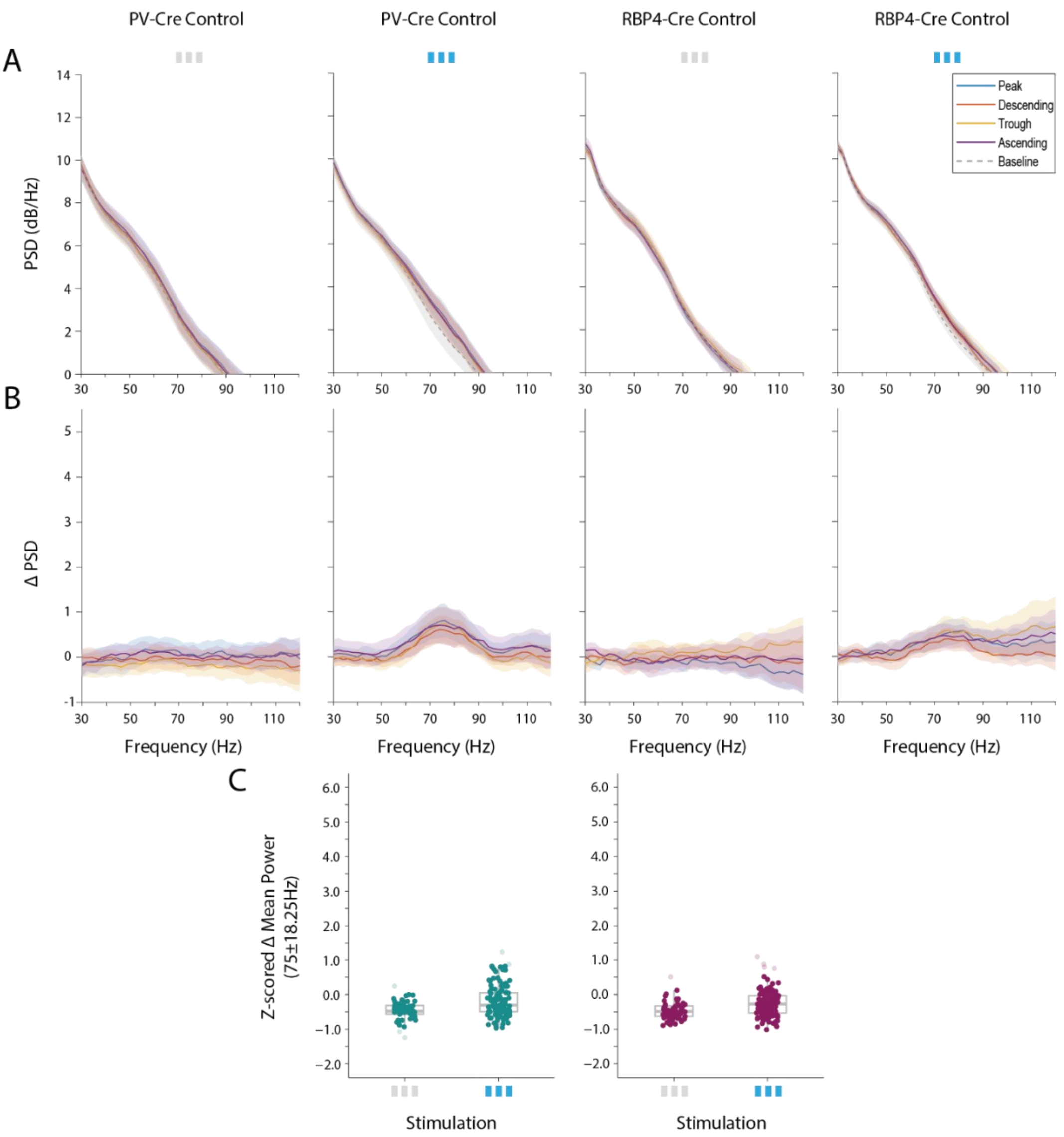


**S7: Closed-loop gamma stimulation of PV+ interneurons and RBP4+ pyramidal neurons led to no phase-dependent effects on local gamma power in GFP-control mice. (A)** Mean power spectral density from 30-120Hz for PV-Cre GFP-control (n=4; 14 recordings/phase stim off & 29-30 recordings/phase stim on) and RBP4-Cre GFP-control (n=4: 15 recordings/phase stim off & 30 recordings/phase stim on) mice, targeting four different phases of the theta oscillation with gamma stimulation either unstimulated (0mW - i.e. using reference time points when stimulation would have been applied) or at 1 or 2mW. Shaded areas show ±2*SEM. **(B)** As in (A) but using baseline-subtracted spectra. **(C)** Statistical analyses performed demonstrated no significant main effect or interaction terms including phase on gamma power (all $p>0.05$; LME), subsequently, data from all phases was combined for further analyses. Boxplots with the z-scored change in mean gamma power with respect to baseline extracted between in the gamma band (56.75-93.25Hz) for PV-Cre and RBP4-Cre GFP-control mice during gamma stimulation (1 or 2mW) vs no stimulation. No significant differences ($p<0.05$) between stimulation on and off were identified in GFP-control mice of either genotype (LME – significant group*stimulation interaction; followed by pairwise post hoc t-tests with Tukey adjustment).

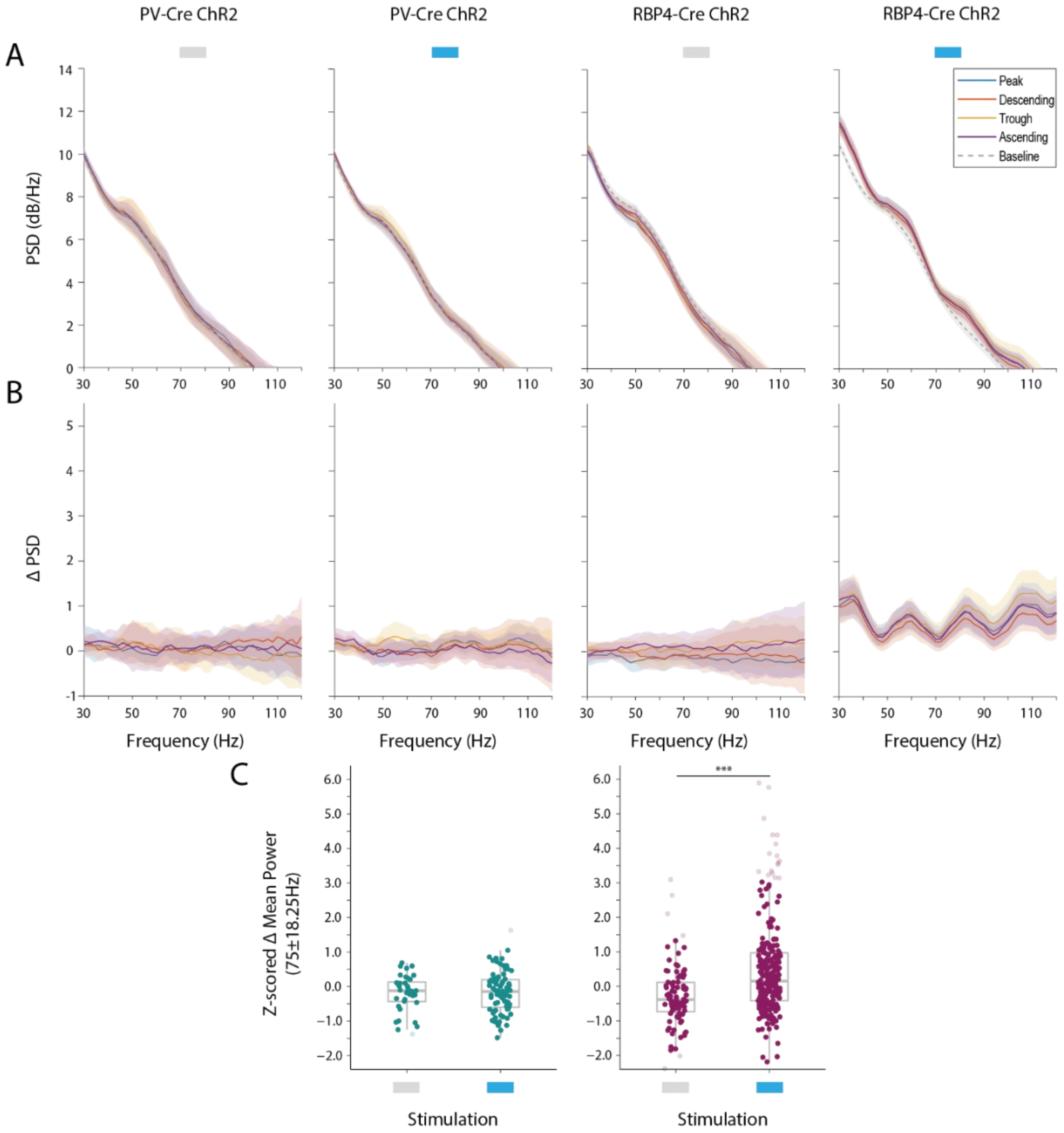


**S8: Closed-loop continuous stimulation led to phase-dependent effects on local gamma power only when targeting RBP4+ pyramidal neurons. (A)** Mean power spectral density from 30-120Hz for PV-Cre ChR2 (n=4; 8-10 recordings/phase stim off & 18 recordings/phase stim on) and RBP4-Cre ChR2 (n=6-10: 21-23 recordings/phase stim off & 60 recordings/phase stim on) mice, targeting four different phases of the theta oscillation with gamma stimulation either unstimulated (0mW - i.e. using reference time points when stimulation would have been applied) or at 1 or 2mW. Shaded areas show ±2*SEM. **(B)** As in (A) but using baseline-subtracted spectra. **(C)** Statistical analyses performed demonstrated no significant main effect or interaction terms including phase on gamma power (all $p>0.05$; LME), subsequently, data from all phases was combined for further analyses. Boxplots with the z-scored change in mean gamma power with respect to baseline extracted between in the gamma band (56.75-93.25Hz) for PV-Cre and RBP4-Cre ChR2 mice during gamma stimulation (1 or 2mW) vs unstimulated. A significant group*stimulation interaction (LME; $F(3, 751)=7.93$, $p<0.001$) was identified, this included data from both GFP-control and ChR2 mice of each genotype. To follow up post hoc t-tests identified significant differences only in RBP4-Cre ChR2 mice ($p<0.001$).

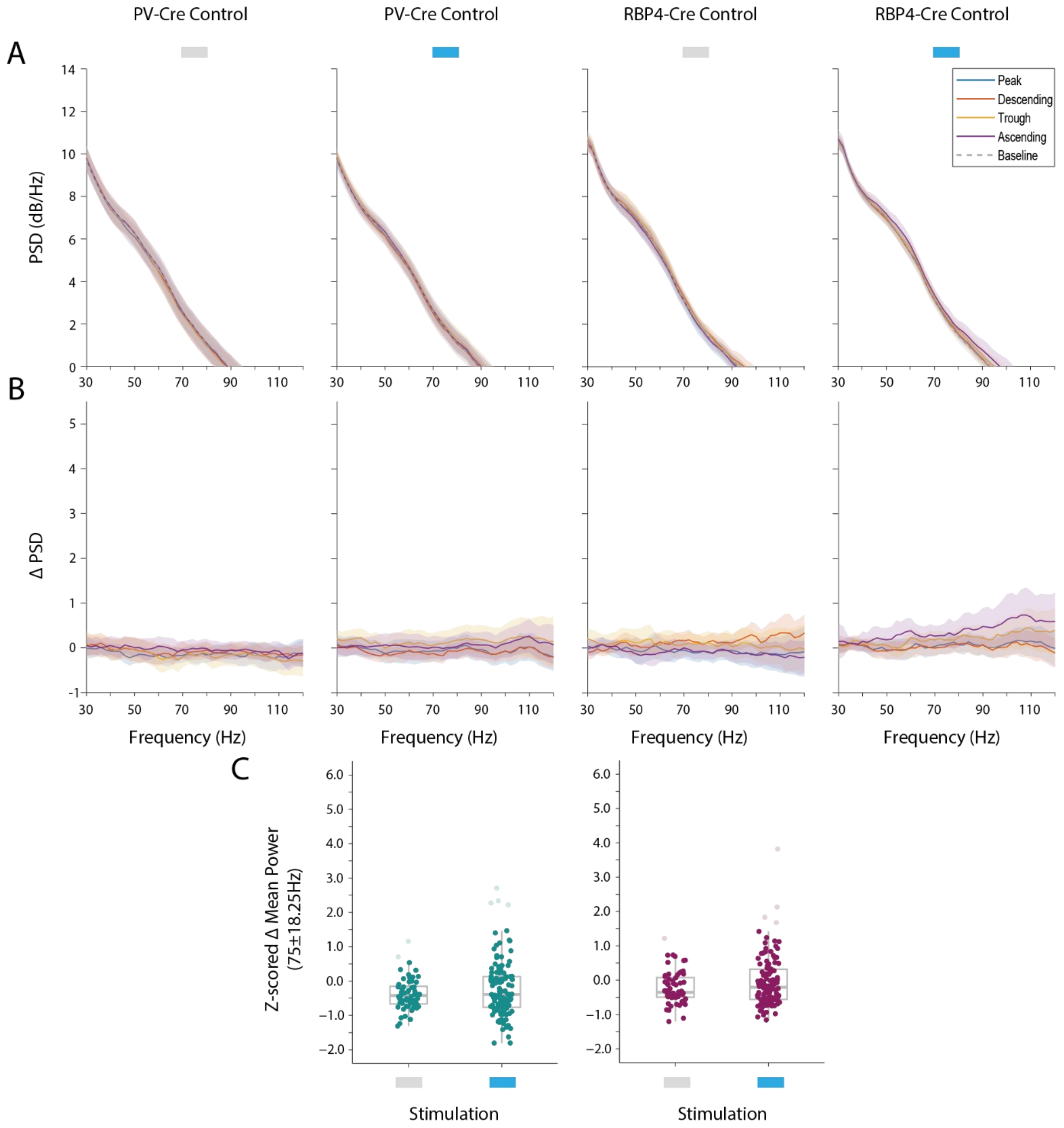


**S9: Closed-loop continuous stimulation of PV+ interneurons and RBP4+ pyramidal neurons led to no phase-dependent effects on local gamma power in GFP-control mice. (A)** Mean power spectral density from 30-120Hz for PV-Cre GFP-control (n=4; 16 recordings/phase stim off & 31 recordings/phase stim on) and RBP4-Cre GFP-control (n=4: 15 recordings/phase stim off & 30 recordings/phase stim on) mice, targeting four different phases of the theta oscillation with continuous stimulation either unstimulated (0mW - i.e. using reference time points when stimulation would have been applied) or at 1 or 2mW. Shaded areas show ±2*SEM. **(B)** As in (A) but using baseline-subtracted spectra. **(C)** Statistical analyses performed demonstrated no significant main effect or interaction terms including phase on gamma power (all $p>0.05$; LME), subsequently, data from all phases was combined for further analyses. Boxplots with the z-scored change in mean gamma power with respect to baseline extracted between in the gamma band (56.75-93.25Hz) for PV-Cre and RBP4-Cre GFP-control mice during continuous stimulation (1 or 2mW) vs unstimulated. No significant differences ($p<0.05$) between stimulation on and off were identified in GFP-control mice of either genotype (LME – significant group*stimulation interaction; followed by pairwise post hoc t-tests with Tukey adjustment).
