## Supplemental-Methods for "Modulation of motor cortical theta and gamma oscillations using phase-targeted, closed-loop optogenetic stimulation of local excitatory and inhibitory neurons"

### Histology - Immunohistochemistry

For post-hoc histology, mice were terminally anaesthetised and transcardially perfused with 4% PFA in 0.1M PBS. The brains were extracted and post-fixed overnight in the same solution at 4°C. Brains were then washed in PBS and stored in PBS-Azide (PBS with 0.05% sodium azide (PBS-Az)). Coronal sections (50µm) were cut into serial wells using a vibrating microtome or freezing sledge microtome. If brains were being frozen, they were suspended in 30% sucrose in 0.1M PB solution for cryoprotection and allowed to descend to the bottom of the vial (typically >24hrs), sections were washed 3x in PBS pre-storage. Sections were then stored in PBS-Az at 4°C until they were used for immunohistochemistry.

Free floating PV-Cre or RBP4-Cre brain sections for immunohistochemistry (1 in 3 sections; from around AP+2.57 to AP -0.49) were washed with gentle shaking in PBS-Triton-Az (PBS-Tx, 0.3% Triton (v/v), 0.02% sodium azide (w/v)) for 3x 10 minutes to permeabilize the tissue. Sections were blocked in 20% v/v normal donkey serum (NDS, Vector Laboratories, California, USA) in PBS-Tx for one hour at room temperature. Following this, sections were incubated in primary antibody solution in PBS-Tx with 1% NDS overnight at room temperature before 3x 10-minute washes in PBS. Primary antibodies were chosen to enhance native green fluorescent protein (GFP) signal alongside delineation of cortical layers, these included GFP (Aves; 1:500), parvalbumin (Synaptic Systems; 1:1000), FoxP2 (Santa Cruz or Abcam; 1:500) and Necab1 (Life Technologies; 1:1000 or Abnova; 1:500). Sections were subsequently incubated using species specific fluorophore-conjugated secondary antibodies (Alexa-488 (1:1000), AMCA (1:250), Alexa-647 (1:250), Cy3 (1:400); Jackson Laboratories) (raised in Donkey) diluted in PBS-Tx and 1% NDS either for 4 hours at room temperature, or overnight at 2-4°C. Following 3 final 10-minute washes in PBS, sections were mounted on glass microscope slides. Coverslips were applied with an aqueous mounting medium (Vectashield, Vector laboratories).

### Microscopy

Images were taken on an Axio Imager M2 epifluorescent microscope (Zeiss), equipped with Plan-Apochromat objective lenses, a Hamamatsu Flash 4.0 LT camera (C10600) and Colibri LEDs. ZEN Blue software was used to acquire tiled single plane images with a 10x objective. A Zeiss LSM 880 Axio Imager 2 laser scanning confocal microscope was used to take higher resolution images. ZEN Black software was used in acquisition of tile-scans using a 5x or 20x Plan-Apochromat air objective lens (NA 0.16 & NA 0.8 respectively). Images were stitched offline using ZEN Blue software. Neuronal specificity images were acquired using a 63x Plan-Apochromat oil objective lens (NA 1.46). AMCA, Alexa-488, Cy3 & Alexa-647 fluorescence were imaged using excitation from a Diode 405nm laser, Argon 488nm laser, and HeNe 543nm laser and HeNe 633nm laser, with MBS-405 and MBS-488/543/633 beamsplitters respectively.
