## Supplemental-Statistical-Tables for "Modulation of motor cortical theta and gamma oscillations using phase-targeted, closed-loop optogenetic stimulation of local excitatory and inhibitory neurons"

### Supplementary Statistic Tables

**Supplementary Table 1: Modulation of theta power following continuous stimulation of PV+ interneurons.**

| Fixed effects | Sum Sq | Mean Sq | NumDF | DenDF | F value | Pr(>F) |
|---|---|---|---|---|---|---|
| Group | 6.66 | 6.66 | 1 | 14.05 | 8.40 | 0.0117 |
| Phase | 2.61 | 0.87 | 3 | 288.21 | 1.10 | 0.3513 |
| Stimulation | 3.82 | 3.82 | 1 | 288.12 | 4.82 | 0.0290 |
| Group*Phase | 11.97 | 3.99 | 3 | 288.21 | 5.03 | 0.0021 |
| Group*Stimulation | 0.07 | 0.07 | 1 | 288.12 | 0.08 | 0.7729 |
| Phase*Stimulation | 1.72 | 0.57 | 3 | 288.21 | 0.72 | 0.5398 |
| Group*Phase*Stimulation | 5.08 | 1.69 | 3 | 288.21 | 2.13 | 0.0961 |

**Supplementary Table 2: Modulation of theta power following continuous stimulation of PV+ interneurons, STIM-ON.**

| Fixed effects | Sum Sq | Mean Sq | NumDF | DenDF | F value | Pr(>F) |
|---|---|---|---|---|---|---|
| Group | 5.06 | 5.06 | 1 | 12.78 | 6.32 | 0.0262 |
| Phase | 3.94 | 1.31 | 3 | 183.66 | 1.64 | 0.1819 |
| Group*Phase | 21.80 | 7.27 | 3 | 183.66 | 9.07 | 1.256e-05 |

**Supplementary Table 3: Modulation of theta power following continuous stimulation of RBP4+ pyramidal neurons.**

| Fixed effects | Sum Sq | Mean Sq | NumDF | DenDF | F value | Pr(>F) |
|---|---|---|---|---|---|---|
| Group | 1.30 | 1.30 | 1 | 14.70 | 2.21 | 0.1581 |
| Phase | 10.56 | 3.52 | 3 | 469.67 | 5.98 | 0.0005 |
| Stimulation | 11.97 | 11.97 | 1 | 478.11 | 20.36 | 8.107e-06 |
| Group*Phase | 6.18 | 2.06 | 3 | 469.67 | 3.50 | 0.0155 |
| Group*Stimulation | 7.14 | 7.14 | 1 | 478.11 | 12.14 | 0.0005 |
| Phase*Stimulation | 1.48 | 0.49 | 3 | 469.67 | 0.84 | 0.4726 |
| Group*Phase*Stimulation | 11.82 | 3.94 | 3 | 469.67 | 6.70 | 0.0002 |

**Supplementary Table 4: Modulation of theta power following continuous stimulation of RBP4+ pyramidal neurons, STIM-ON.**

| Fixed effects | Sum Sq | Mean Sq | NumDF | DenDF | F value | Pr(>F) |
|---|---|---|---|---|---|---|
| Group | 3.01 | 3.01 | 1 | 16.05 | 4.88 | 0.0419 |
| Phase | 15.89 | 5.30 | 3 | 323.26 | 8.58 | 1.69e-05 |
| Group*Phase | 27.64 | 9.21 | 3 | 323.26 | 14.93 | 3.99e-09 |

**Supplementary Table 5: Modulation of theta power following gamma frequency stimulation of PV+ interneurons.**

| Fixed effects | Sum Sq | Mean Sq | NumDF | DenDF | F value | Pr(>F) |
|---|---|---|---|---|---|---|
| Group | 0.01 | 0.01 | 1 | 17.41 | 0.02 | 0.9035 |
| Phase | 0.52 | 0.17 | 3 | 324.02 | 0.26 | 0.8570 |
| Stimulation | 1.44 | 1.44 | 1 | 325.28 | 2.14 | 0.1448 |
| Group*Phase | 8.66 | 2.89 | 3 | 324.02 | 4.29 | 0.0055 |
| Group*Stimulation | 0.16 | 0.16 | 1 | 325.28 | 0.24 | 0.6244 |
| Phase*Stimulation | 7.34 | 2.45 | 3 | 323.76 | 3.63 | 0.0133 |
| Group*Phase*Stimulation | 1.86 | 0.62 | 3 | 323.76 | 0.92 | 0.4304 |

**Supplementary Table 6: Modulation of theta power following gamma frequency stimulation of PV+ interneurons, STIM-ON.**

| Fixed effects | Sum Sq | Mean Sq | NumDF | DenDF | F value | Pr(>F) |
|---|---|---|---|---|---|---|
| Group | 0.00 | 0.00 | 1 | 19.36 | 0.00 | 0.9725 |
| Phase | 7.01 | 2.34 | 3 | 205.18 | 3.85 | 0.0105 |
| Group*Phase | 11.51 | 3.84 | 3 | 205.18 | 6.32 | 0.0004 |

**Supplementary Table 7: Modulation of theta power following gamma frequency stimulation of RBP4+ pyramidal neurons.**

| Fixed effects | Sum Sq | Mean Sq | NumDF | DenDF | F value | Pr(>F) |
|---|---|---|---|---|---|---|
| Group | 1.06 | 1.06 | 1 | 18.12 | 1.87 | 0.1887 |
| Phase | 6.43 | 2.14 | 3 | 544.31 | 3.78 | 0.0105 |
| Stimulation | 8.66 | 8.66 | 1 | 545.33 | 15.27 | 0.0001 |
| Group*Phase | 4.50 | 1.50 | 3 | 544.31 | 2.65 | 0.0484 |
| Group*Stimulation | 14.44 | 14.44 | 1 | 545.33 | 25.46 | 6.172e-07 |
| Phase*Stimulation | 5.73 | 1.91 | 3 | 544.31 | 3.36 | 0.0185 |
| Group*Phase*Stimulation | 4.32 | 1.44 | 3 | 544.31 | 2.54 | 0.0560 |

**Supplementary Table 8: Modulation of theta power following gamma frequency stimulation of RBP4+ pyramidal neurons, STIM-ON.**

| Fixed effects | Sum Sq | Mean Sq | NumDF | DenDF | F value | Pr(>F) |
|---|---|---|---|---|---|---|
| Group | 2.53 | 2.53 | 1 | 17.17 | 4.05 | 0.0600 |
| Phase | 14.25 | 4.75 | 3 | 324.34 | 7.63 | 6.1e-05 |
| Group*Phase | 10.88 | 3.63 | 3 | 324.34 | 5.82 | 0.0007 |

**Supplementary Table 9: Gamma rhythmicity does not alter phase-dependent theta power (stimulation pattern: gamma, arrhythmic gamma).**

| Fixed effects | Sum Sq | Mean Sq | NumDF | DenDF | F value | Pr(>F) |
|---|---|---|---|---|---|---|
| Stimulation pattern | 3.79 | 3.80 | 1 | 300.24 | 7.89 | 0.0053 |
| Stimulation | 29.24 | 29.24 | 1 | 300.01 | 60.88 | 1.017e-13 |
| Phase | 23.36 | 7.79 | 3 | 299.00 | 16.21 | 8.720e-10 |
| Stimulation pattern*Stimulation | 0.01 | 0.01 | 1 | 300.24 | 0.02 | 0.9016 |
| Stimulation pattern*Phase | 0.88 | 0.30 | 3 | 299.00 | 0.61 | 0.6095 |
| Stimulation*Phase | 22.02 | 7.34 | 3 | 299.00 | 15.28 | 2.828e-09 |
| Stimulation pattern*Stimulation*Phase | 0.44 | 0.15 | 3 | 299.00 | 0.31 | 0.8211 |

**Supplementary Table 10: Stimulation pattern does not affect phase-dependent theta power modulation, STIM-ON (Stimulation pattern: gamma, arrhythmic gamma, continuous).**

| Fixed effects | Sum Sq | Mean Sq | NumDF | DenDF | F value | Pr(>F) |
|---|---|---|---|---|---|---|
| Stimulation pattern | 3.73 | 1.86 | 2 | 277.92 | 2.73 | 0.0676 |
| Phase | 81.83 | 27.28 | 3 | 277.92 | 39.92 | <2e-16 |
| Stimulation pattern*Phase | 4.50 | 0.75 | 6 | 277.92 | 1.10 | 0.3641 |

**Supplementary Table 11: No effect of phase on the gamma band following gamma frequency stimulation of PV+ interneurons.**

| Fixed effects | Sum Sq | Mean Sq | NumDF | DenDF | F value | Pr(>F) |
|---|---|---|---|---|---|---|
| Group | 210 | 210 | 1 | 15.06 | 0.23 | 0.64 |
| Phase | 490 | 163 | 3 | 314.16 | 0.17 | 0.91 |
| Stimulation | 32213 | 32213 | 1 | 316.00 | 34.30 | 1.184e-08 |
| Group*Phase | 1506 | 502 | 3 | 314.16 | 0.54 | 0.66 |
| Group*Stimulation | 290 | 290 | 1 | 316.00 | 0.31 | 0.58 |
| Phase*Stimulation | 844 | 281 | 3 | 313.81 | 0.30 | 0.83 |
| Group*Phase*Stimulation | 87 | 29 | 3 | 313.81 | 0.03 | 0.99 |

**Supplementary Table 12: No effect of phase on the gamma band following continuous stimulation of PV+ interneurons.**

| Fixed effects | Sum Sq | Mean Sq | NumDF | DenDF | F value | Pr(>F) |
|---|---|---|---|---|---|---|
| Group | 459.40 | 459.40 | 1 | 13.07 | 1.11 | 0.31 |
| Phase | 193.88 | 64.63 | 3 | 273.94 | 0.16 | 0.93 |
| Stimulation | 612.80 | 612.80 | 1 | 273.81 | 1.48 | 0.23 |
| Group*Phase | 319.05 | 106.35 | 3 | 273.94 | 0.26 | 0.86 |
| Group*Stimulation | 198.47 | 198.47 | 1 | 273.81 | 0.48 | 0.49 |
| Phase*Stimulation | 865.95 | 288.65 | 3 | 273.94 | 0.70 | 0.56 |
| Group*Phase*Stimulation | 50.09 | 16.70 | 3 | 273.94 | 0.04 | 0.99 |

**Supplementary Table 13: No effect of phase on the gamma band following gamma frequency stimulation of RBP4+ pyramidal neurons.**

| Fixed effects | Sum Sq | Mean Sq | NumDF | DenDF | F value | Pr(>F) |
|---|---|---|---|---|---|---|
| Group | 10576 | 10576 | 1 | 17.62 | 3.96 | 0.0622 |
| Phase | 1120 | 373 | 3 | 546.50 | 0.14 | 0.9361 |
| Stimulation | 341347 | 341347 | 1 | 548.67 | 127.93 | <2e-16 |
| Group*Phase | 799 | 266 | 3 | 546.50 | 0.10 | 0.9601 |
| Group*Stimulation | 187188 | 187188 | 1 | 548.67 | 70.16 | 4.6e-16 |
| Phase*Stimulation | 1277 | 426 | 3 | 546.50 | 0.16 | 0.9235 |
| Group*Phase*Stimulation | 1396 | 465 | 3 | 546.50 | 0.17 | 0.9137 |

**Supplementary Table 14: No effect of phase on the gamma band following continuous stimulation of RBP4+ pyramidal neurons.**

| Fixed effects | Sum Sq | Mean Sq | NumDF | DenDF | F value | Pr(>F) |
|---|---|---|---|---|---|---|
| Group | 592.6 | 592.6 | 1 | 14.00 | 0.57 | 0.4644 |
| Phase | 2628.9 | 876.3 | 3 | 471.22 | 0.84 | 0.4742 |
| Stimulation | 24174.7 | 24174.7 | 1 | 485.63 | 23.08 | 2.072e-06 |
| Group*Phase | 330.2 | 110.1 | 3 | 471.22 | 0.11 | 0.9571 |
| Group*Stimulation | 10434.5 | 10434.5 | 1 | 485.63 | 9.96 | 0.0017 |
| Phase*Stimulation | 1498.0 | 499.3 | 3 | 471.22 | 0.48 | 0.6986 |
| Group*Phase*Stimulation | 2165.4 | 721.8 | 3 | 471.22 | 0.69 | 0.5590 |

**Supplementary Table 15: Modulation of gamma power following gamma frequency stimulation of PV+ interneurons & RBP4+ pyramidal neurons.**

| Fixed effects | Sum Sq | Mean Sq | NumDF | DenDF | F value | Pr(>F) |
|---|---|---|---|---|---|---|
| Group | 4.20 | 1.40 | 3 | 35.29 | 3.07 | 0.040 |
| Stimulation | 57.48 | 57.48 | 1 | 860.64 | 125.96 | <2e-16 |
| Group*Stimulation | 68.64 | 22.88 | 3 | 860.78 | 50.14 | <2e-16 |

**Supplementary Table 16: Modulation of gamma power following continuous stimulation of PV+ interneurons & RBP4+ pyramidal neurons.**

| Fixed effects | Sum Sq | Mean Sq | NumDF | DenDF | F value | Pr(>F) |
|---|---|---|---|---|---|---|
| Group | 1.54 | 0.51 | 3 | 26.82 | 0.69 | 0.5674 |
| Stimulation | 12.48 | 12.48 | 1 | 749.11 | 16.77 | 4.687e-05 |
| Group*Stimulation | 17.71 | 5.91 | 3 | 751.04 | 7.93 | 3.27e-05 |

**Supplementary Table 17: Open-loop replay of gamma frequency stimulation modulates gamma power to a lesser extent than closed-loop stimulation in RBP4-Cre mice.**

| Fixed effects | Sum Sq | Mean Sq | NumDF | DenDF | F value | Pr(>F) |
|---|---|---|---|---|---|---|
| Group | 1.01 | 1.01 | 1 | 16.15 | 3.63 | 0.0746 |
| Stimulation type | 5.24 | 5.24 | 1 | 366.00 | 18.82 | 1.859e-05 |
| Group*Stimulation type | 3.90 | 3.90 | 1 | 366.00 | 14.03 | 0.0002 |
